# ThermoPlex: An Automated Design Tool for Target-specific Multiplex PCR Primers based on DNA Thermodynamics

**DOI:** 10.1101/2024.12.27.630477

**Authors:** Altair Agmata, Kevin Labrador, Joseph Dominic Palermo, Maria Josefa Pante

## Abstract

Multiplex PCR-based assays are indispensable platforms for rapid and cost-effective DNA-based multi-target detection. The success of such an assay highly depends on the accurate design of oligonucleotide primers, arguably its most vital component. In this study, the ThermoPlex design tool is introduced, offering an automated design pipeline for target-specific multiplex PCR primers motivated by DNA thermodynamics. From a sequence alignment of all relevant target and non-target sequences, ThermoPlex automatically designs multiplex PCR primer candidates in just a matter of minutes. The software also offers tools for thermodynamic calculations that can either be used apart from the automated primer screening routine or in conjunction with other existing primer design tools, depending on the on the needs of the user. Evidences presented here in this study provide insights on the performance of the software through theoretical and experimental analyses, serving to establish the reliability of its framework.

## 1 Introduction

Multiplex PCR-based assays remain widely employed for rapid DNA-based target detection due to its multi-target detection capabilities in a single assay platform [1]. Through the years, various detection and identification assays have emerged, utilizing this platform for a wide variety of organisms, including microbial [2, 3], plant [4], and animal [5, 6, 7, 8, 9] species. Recently, with the emergence of the coronavirus disease 2019 (COVID-19), numerous multiplex PCR assays for clinical diagnostics have been developed commercially and publicly [10, 11, 12, 13] taking advantage of the assay’s ability to simultaneously detect the target along with multiple non-target samples in a single PCR reaction system. Multiplex PCR assays are thus expected to remain a relevant assay platform in the subsequent years, owing to their versatility in a variety of fields of application, as well as their cost-effectiveness.

The success of a PCR assay heavily depends on the design of its primers, perhaps its most vital component. In fact, PCR assay design issues typically arise from the designer’s lack of experience with the primer design process [14]. This includes understanding of key parameters to optimize and familiarity with appropriate design tools to generate optimal primers. The complexity of the problem further escalates when trying to design a multiplex PCR assay. Apart from the primer’s interaction with the target DNA, one must consider the interaction of multiple primers among themselves as all these components interact together in a single PCR solution system. These interactions result to numerous undesirable hybridization reactions affecting the assay’s overall specificity and efficiency. Tackling such complexity with rigor, researchers often resort to an iterative, expensive and labor-intensive workflow [15] with high incidence of failure even after extensive experimentation [16].

The design challenges described can be alleviated by in silico approaches that systematically evaluate the performance of primers before undergoing experimental scrutiny. The most common type of in silico method for primer design evaluate primer performance through heuristics driven by practical experience with PCR (e.g., melting point matching, base pair mismatches, BLAST/alignment scores). However some of the major caveats of these approaches include the lack of direct physical interpretation, and reliance on parameters with arbitrarily chosen thresholds [17]. Such approaches can thus be considered lacking in terms of proper design of multiplex PCR assays. In contrast, approaches based on nucleic acid thermodynamics can overcome these limitations. Thermodynamics of nucleic acid interactions have been extensively studied in the past two decades, with values for useful thermodynamic parameters (enthalpy (Δ*H*), entropy (Δ*S*), and Gibbs free energy (Δ*G*) changes) already been compiled for every possible interaction motifs [18, 19, 20, 21, 22, 23]. This makes it possible to quantitatively predict thermodynamic interactions between nucleic acids of any base pair sequence. Moreover, the use of thermodynamic parameters through the Nearest-Neighbor (NN) [24, 25] model have been proven to yield fairly accurate results consistent to that of experimental observations for oligonucleotide interactions [18]. This makes nucleic acid thermodynamics a solid foundation for developing a rigorous approach to designing multiplex PCR primers.

In this study, a design tool called ThermoPlex is introduced, with the aim of assisting with the design and screening of multiplex-compatible, target-specific PCR Primers from target and non-target sequence alignment datasets. Although numerous programs already exist predicting thermodynamic interactions of nucleic acids, very few open-access software focuses on automated and streamlined screening of multiplex PCR primer candidates from a given sequence dataset. The methodology utilizes a novel *O*(*M* × *N* ) time algorithm based on the NN model doublet parameters to predict the ensemble thermodynamics of DNA-DNA interactions. The thermodynamic information is then used to negatively select target-specific primer candidates against undesirable target sequences. The target-specific primer candidates are then grouped together based on binding region overlaps and then assessed for multiplex-compatibility through an algorithm that computationally simulates multi-reaction equilibrium thermodynamics. The ThermoPlex software was designed in MATLAB with user interaction available either through a standalone GUI-enabled app (Windows/MacOS) or through the use of the source code. The standalone app together with the source codes are available at https://github.com/aagmata/ThermoPlex.

## 2 Methodology

### 2.1 *ThermoDHyb*: Thermodynamics of DNA hybridization prediction algorithm

The *ThermoDHyb* algorithm aims to predict the thermodynamics of double-stranded ensemble interaction between two single-stranded DNA. The novel algorithm calculates and predicts the associated Gibbs free energy change (Δ*G*) between a single-stranded reference state and each double-stranded microstate corresponding to local minima in the cumulative pairwise energy matrix *P*. The routine derives its main working principle from the Smith-Waterman algorithm [26] for sequence alignment. Sequence pairwise comparison (Figure 2A) is done per sequence doublet according to the NN model, with values of the P matrix populated according to the equation:

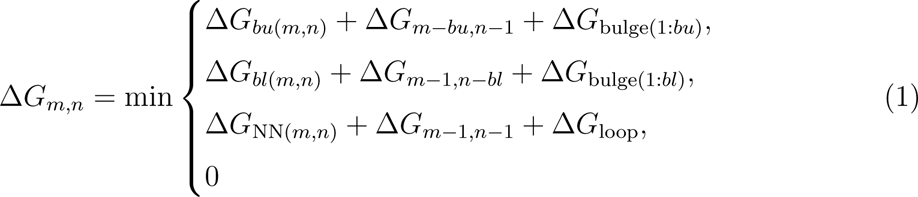

**Figure 1:**
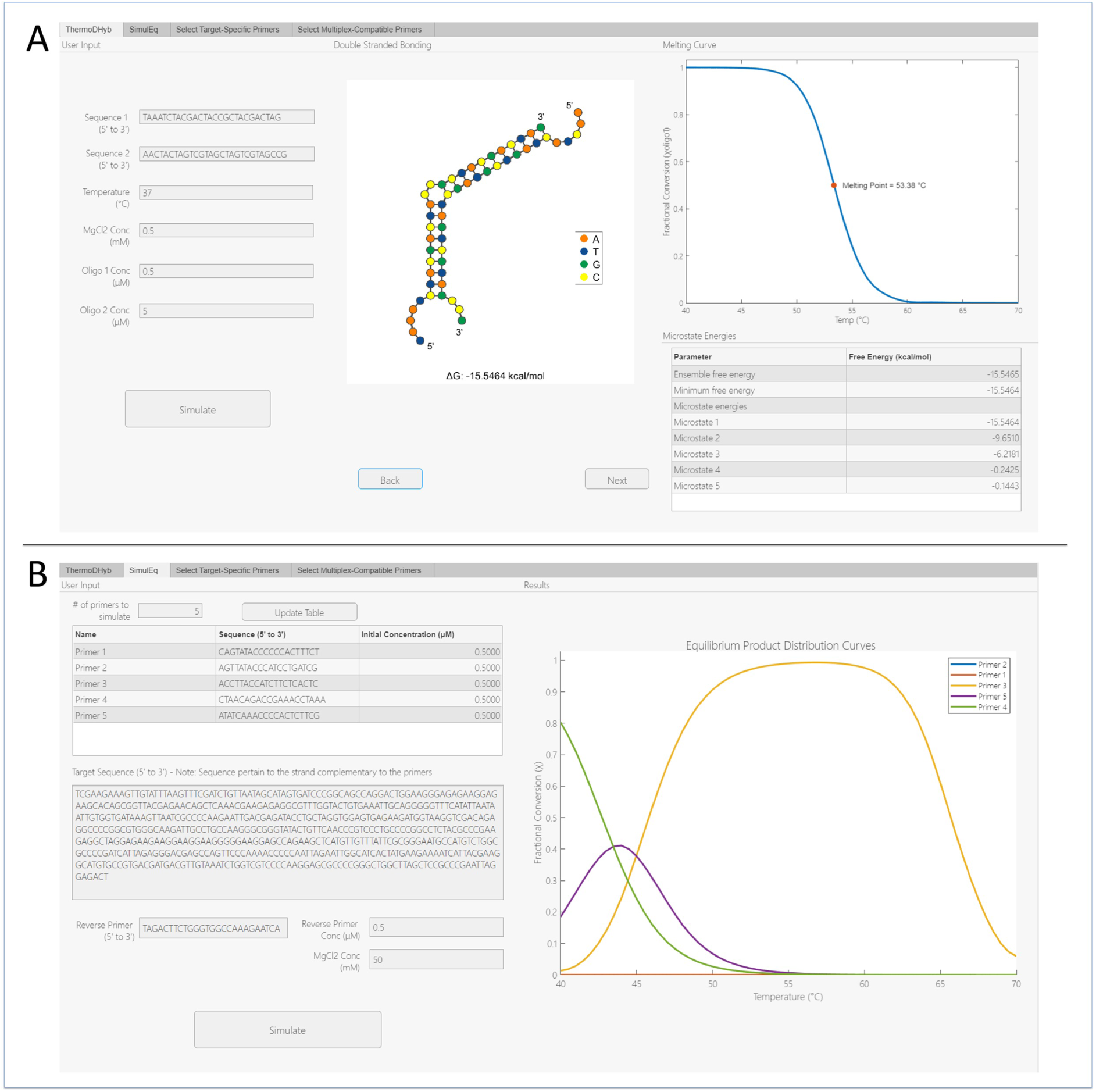
Snapshot of the ThemoPlex software user interface (UI) running the A) ThermoDHyb and B) SimulEq routines.

**Figure 2:**
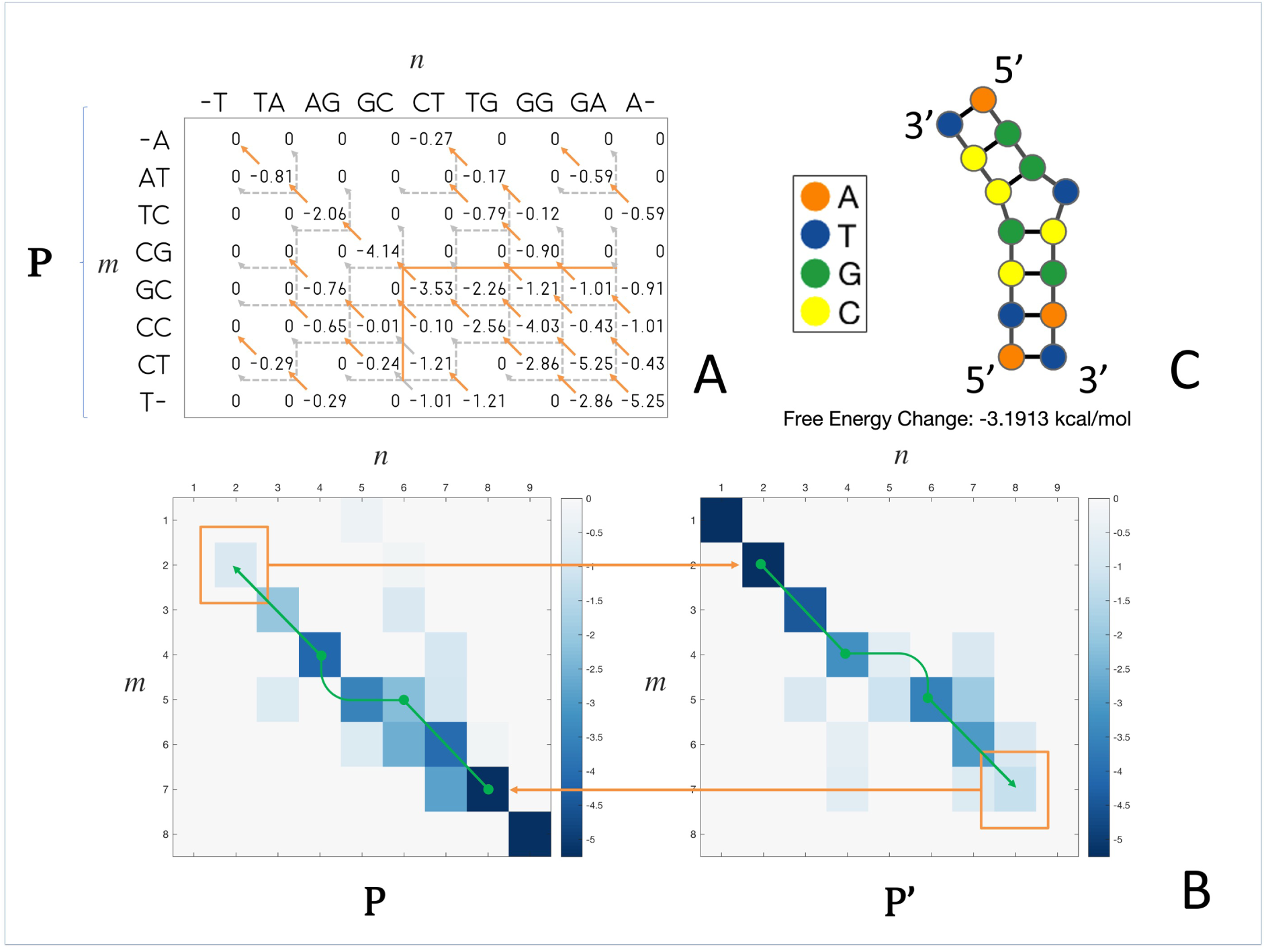
A) Cumulative pairwise energy matrix of doublet pairs for sequences 5’ ATCGCCT 3’ (Sequence *i*) and 5’ AGGTCGAT 3’ (Sequence j) at 40^◦^C and 1M NaCl simulation conditions. Sequence *i* and *j* doublets are listed per row and column respectively. Arrows represent the cumulative dependence of each element (*m*, *n*) to (*m* − bulge, *n* − 1), (*m* − 1*, n* − bulge), or (*m* − 1*, n* − 1) elements. Orange, solid arrows depict the dependence case chosen corresponding to the minimum score case evaluated through equation 1. B) Propagation of cumulative energy scores (down-right for *P* , up-left for *P* ^′^). Terminal doublet pairs are boxed in orange, green arrows show the path of tracing the doublets to infer the minimum free energy (MFE) structure of each microstate. C) Global MFE structure and corresponding free energy change 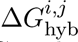 of the most stable microstate predicted by *ThermoDHyb* for sequences *i* and *j* at 40^◦^C and 1M NaCl simulation conditions.) *ThermoDHyb* and B) SimulEq routines.

where Δ*G_NN_* is the cumulative free energy value, while Δ*G_bulge_* and Δ*G_loop_* correspond to length dependent penalties from bulge and internal loop formation, respectively. Δ*G_NN_* values were calculated from enthalpy and entropy values while also accounting for the salt concentration dependence according to Owczarzy’s model [27].

The matrix-filling procedure is performed bi-directionally (Figure 2B) to predict multiple initiation and propagation paths across each local minimum, corresponding to each system microstate *s*. The energies of the set of microstates *σ* are then summed according to the partition function *Z_i,j_*(eq 2) which is in turn used to find the hybridization Gibbs free energy 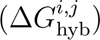 according to equation 3:

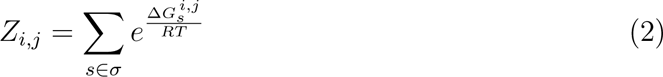

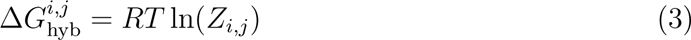

This allows *ThermoDHyb* to calculate thermodynamic interactions that account for significant contributors to the partition function weighed according to the Gibbs factor. Subsequently, the routine traces back the propagation path from the terminal state that along each local minimum to predict the minimum free energy (MFE) structure of each microstate (Figure 2C). The inner workings of the algorithm, together with assumptions made, are explained further in the Supplementary Material – Section 1.

### 2.2 *SiMulEq* : Simulation of Multi-reaction Equilibria interaction

The *SiMulEq* algorithm simulates the concentration of the double-stranded DNA products in thermodynamic equilibrium formed by the primers binding specifically and non-specifically to the given DNA template. The equilibrium scenario is modeled as a set of interdependent competitive reactions that seeks to minimize the total Gibbs free energy of the system. A fully-defined system of equation was derived from the minimization of the total Gibbs free energy as a function of all the species’ chemical potential through the summability rule of partial properties [28]. The function was then constrained with material balance equations representing nucleotide strand conservation. By employing the method of Lagrange multipliers [29], the problem is converted from an optimization problem to a multi-dimensional root-finding problem of the form (derivation at Supplementary Material – Section 2):

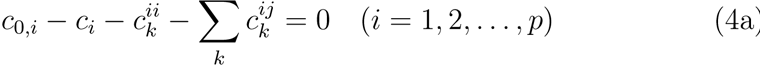

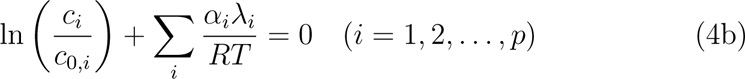

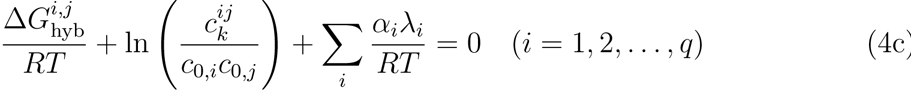

where *c*_0_*,_i_*, *c_i_*, 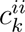 are initial single-stranded, equilibrium single- and double-stranded DNA concenrations in the multiplex solution in standard mol/L, respectively, *α_i_* is a repetition multiplier, *λ_i_*is the lagrange multiplier for each strand species *i* and *j*. The resulting system of equations with 2*p* + *q* equations and unknowns are solved through Newton’s method with finite-difference Jacobian. This yields concentration values of each chemical species at chemical equilibrium for any given temperature. The concentration values are then corrected (see Supplementary Material – Section 3) based on amplification inhibition due to non-target primers binding between the forward target primer and the reverse primer (Figure 4). *SiMulEq* then iterates this procedure for a range of temperatures (40 – 70 ^◦^C) with 2 ^◦^C increments. The final output of *SiMulEq* are equilibrium product distribution (EPD) curves as functions of temperature.

**Figure 3:**
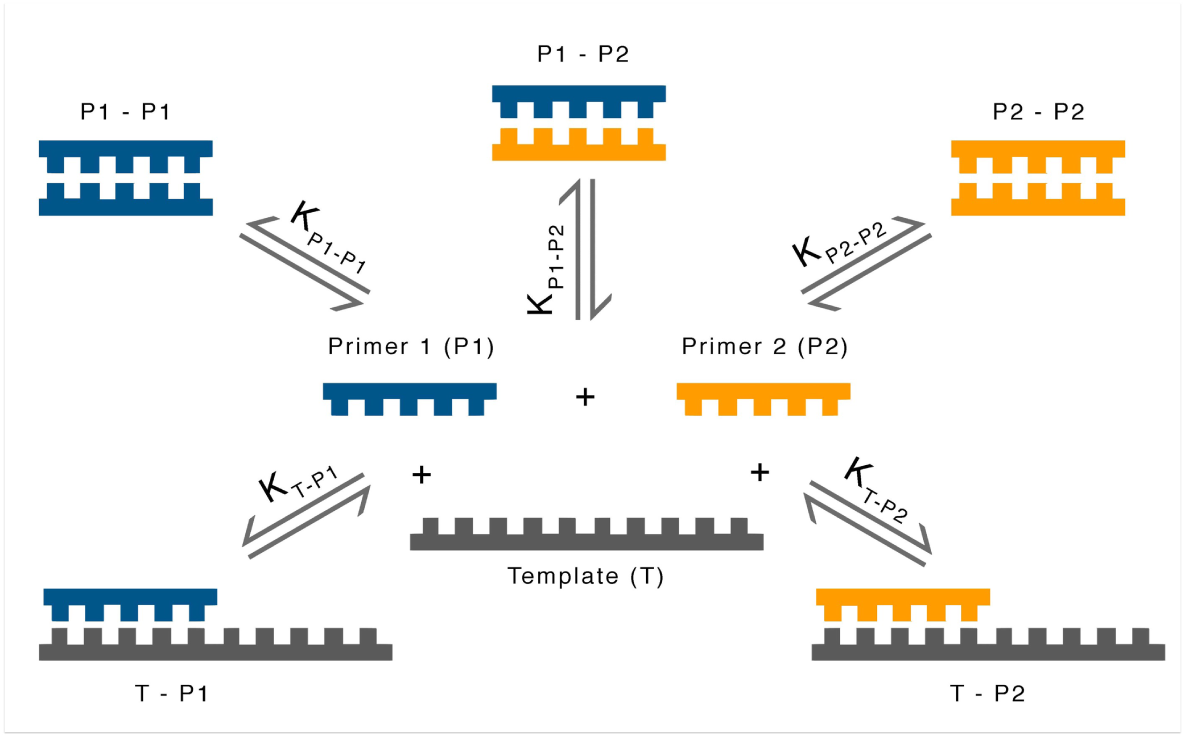
Complex multi-reaction equilibrium scenario accounting for each of the possible pairwise interactions in a two-primer, one-template PCR reaction system. Template-template interactions are not accounted for as it occurs on a different mechanism in the PCR annealing step.

**Figure 4:**
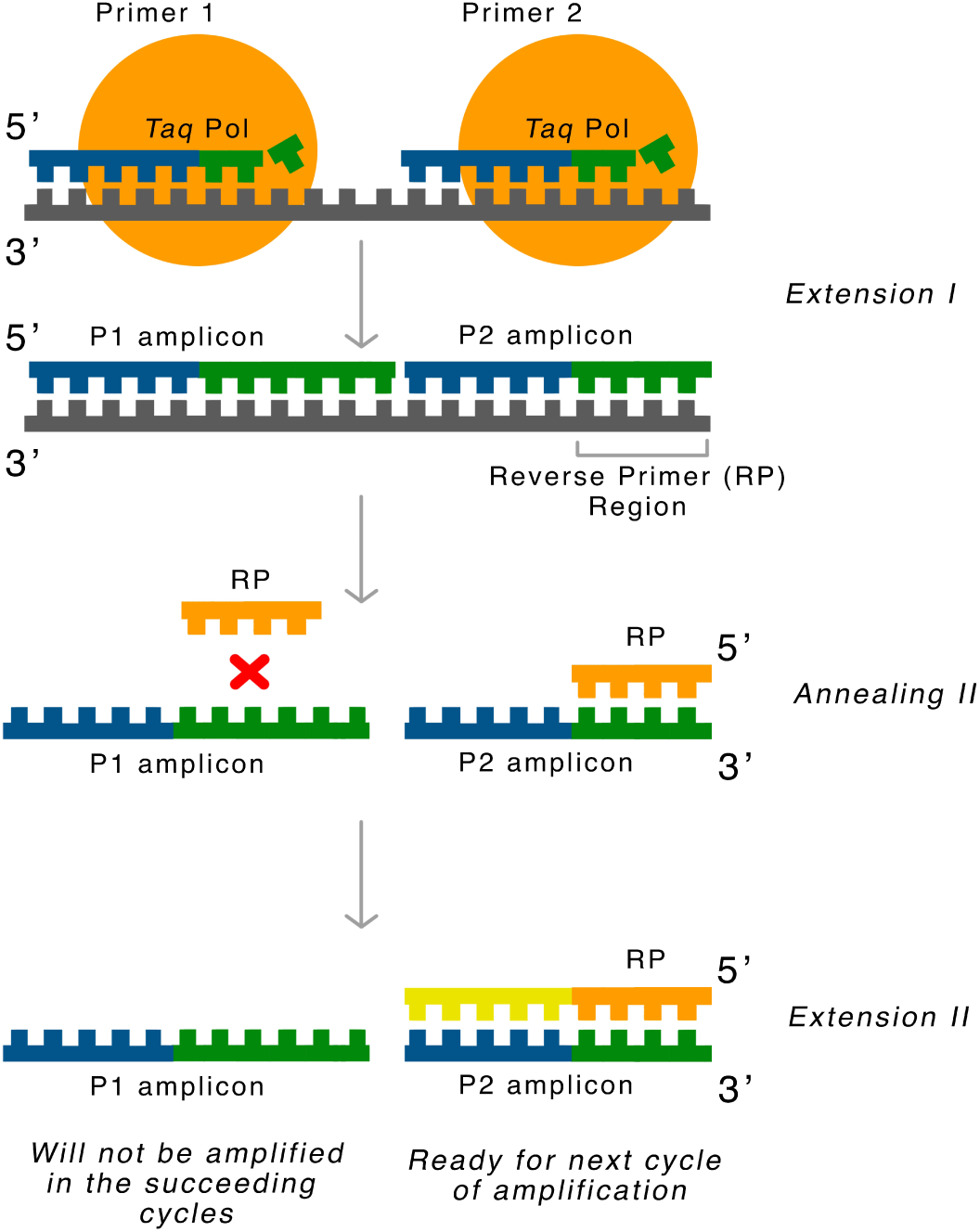
Extension inhibition mechanism where amplification of P1 products is inhibited by a primer (P2) binding along its extension path.

### 2.3 *ThermoPlex* primer selection routine

The ThermoPlex primer selection routine (Figure 5) aims to select multiplex-compatible, target-specific PCR primers from an input alignment of target sequences. Target-specific primers are first selected by iteratively evaluating every possible primer sequence from the consensus target sequence against all the non-target sequence through a moving frame with size equal to the primer length + 2 (the value 2 corresponds to single nucleotide flanking region at 5’ and 3’ end). This assumes that the most favorable interaction with a non-target template of a primer is located on the same region where the sequence was derived from the target sequence. In each iteration, a primer is first evaluated across a set of heuristic criteria (Figure 5*) prior to thermodynamic calculations. Subsequently, calculation of the primers 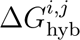 on a single user-specified temperature (arbitrarily around PCR annealing temperature) with the sequence frame of non-target sequence *j* is performed through *ThermoDHyb*. Template fractional conversion *χ_t_*, the mole fraction of the target or non-target template hybridized by the primer, is then calculated according to the equation:

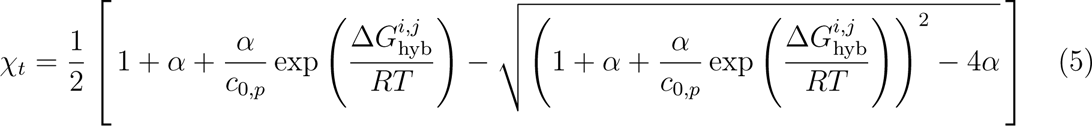

**Figure 5:**
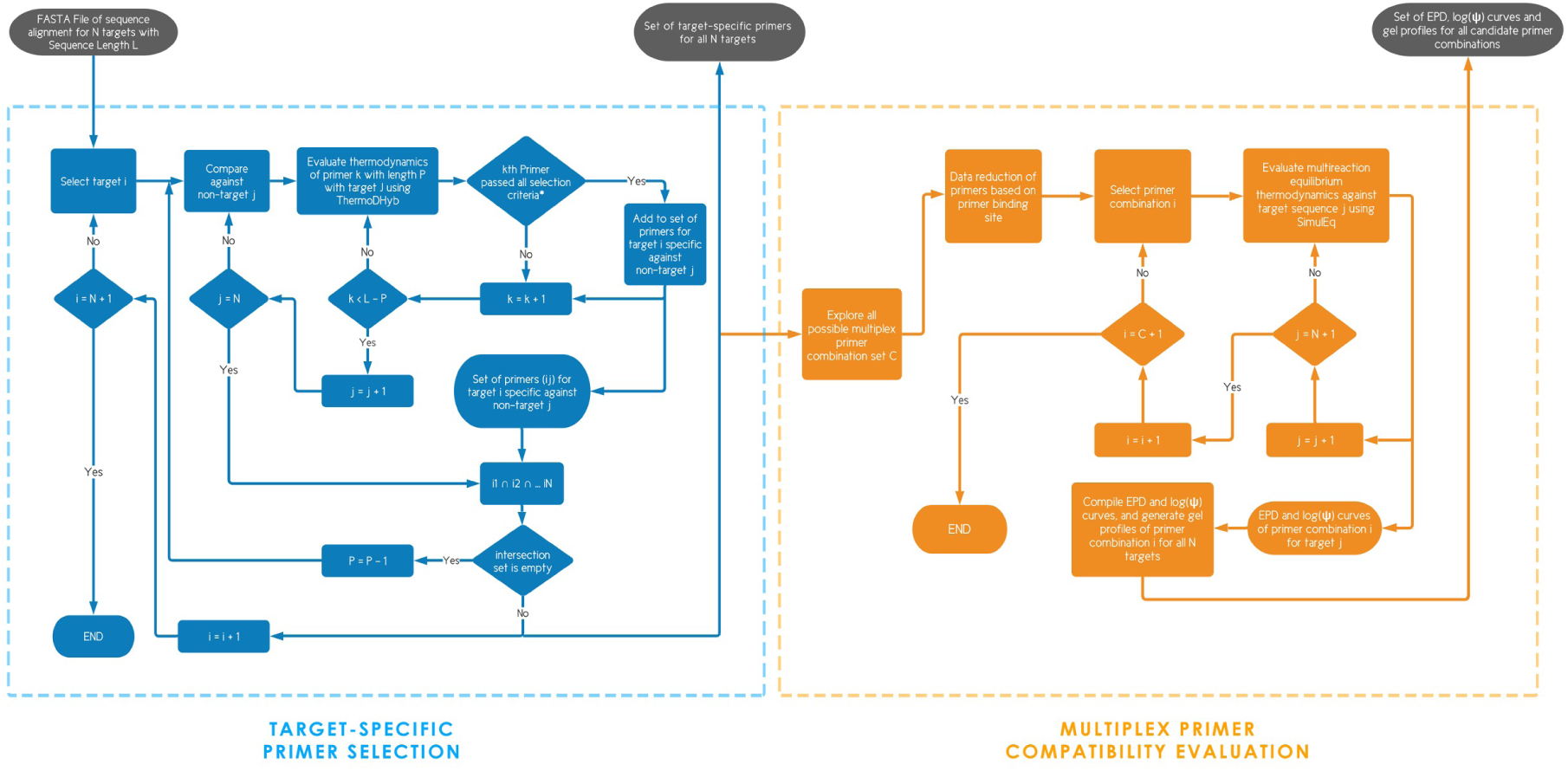
The *ThermoPlex* algorithm subdivided into the target-specific primer selection and multiplex primer compatibility evaluation. *Selection criteria include: no. of mismatch < user-defined mismatch parameter (*M* ); primer *k* does not contain ambiguous nucleotide characters; primer k does not form stable secondary structures (checked through MATLAB’s bioinformatics toolbox function rnafold); fractional specificity (*ψ*) and target fractional conversion 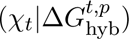 ≥ user-defined threshold.

where:

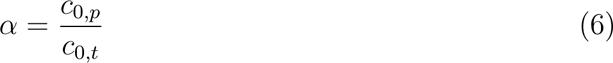

where *c*_0_*_,t_* and *c*_0_*_,p_*, are the initial concentrations of the target template and the primer, respectively. Derivation is explained further in the Supplementary Material (Section 4). The derived model contains the variable *c*_0_*_,t_* in *α*, which is a huge source of uncertainty due to the difficulty of knowing its exact value in terms of molar units. To ensure the robustness of the model despite this uncertainty, a sensitivity analysis for all the relevant variables (*T* , MgCl2 concentration, and *α*) was performed to check whether the uncertainties in *α* is relevant enough. This was implemented through a Monte-Carlo simulation with uniform distribution of input variables.

Fractional specificity (*ψ*) is then calculated as the ratio of *χ_t_* between target (*T* ) to non-target (*N* ):

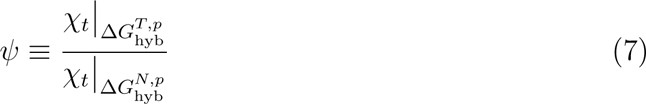

*ψ* describes how probable a primer will hybridize with the target compared to the undesirable non-target sequence (e.g. *ψ* = 1000 means there is 1000 times more primer-target product compared to primer-non-target product upon annealing at the specified temperature). Both *ψ* and 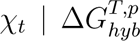 are used as selection criteria based on a user-defined threshold to screen for primer specificity and efficiency, respectively. Primers that passed the criteria are then included in the set ij corresponding to primers derived from target *i* specific against non-target *j*. Upon completion of iteration through all the non-target sequences, the intersection of sets *i*_1_, *i*_2_, . . . *i_N_* (where N is the total number of targets) is then taken, comprising the set of primers for target *i* specific to all the non-target sequences. In the event that the intersection results to an empty set, the process is repeated for a reduced primer length to increase the relative effects of mismatches and thus decrease 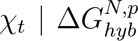 , increasing primer specificity. The whole process is repeated until each target has their own set of target-specific primers.

The set of all target-specific primers for each target are then used to build all possible primer combinations with one primer each target, for a total of *N* primers per combination. The combinations are evaluated on the constraint of amplicon site resolution (*R*). Specifically, each primer in a multiplex combination should be spaced at least *R* bp apart. This will ensure that primers will produce distinguishable gel bands corresponding to each of the target in a multiplex set. Each primer combinations are then evaluated on their multi-reaction equilibrium thermodynamics using *SimulEq*. New values for fractional specificity are also calculated in the context of the multiplex reaction system (*ψ_i_*(*T* )) for each primer *i* corresponding to the desired highest yielding primer, across each temperature increment *T* :

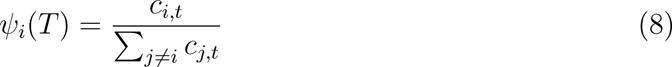

This results to a curve describing the fractional conversion of target *t* from its hybridization with primer *i* relative to all the undesired double-stranded products l due to hybridization between primers *j* and target *t*. The routine will ultimately output EPD curves, *log*(*ψ*) curves and gel profiles for each multiplex primer combination which gives insights on the performance of each multiplex primer set across varying temperature conditions. The ThermoPlex’s subroutines (“Select Target-Specific Primers” and “Select Multiplex-Compatible Primers”) comprising the whole primer selection process were benchmarked in terms of runtimes, with input parameters listed in Table 1, to test how fast each performs across various personal computing platforms.

**Table 1:**
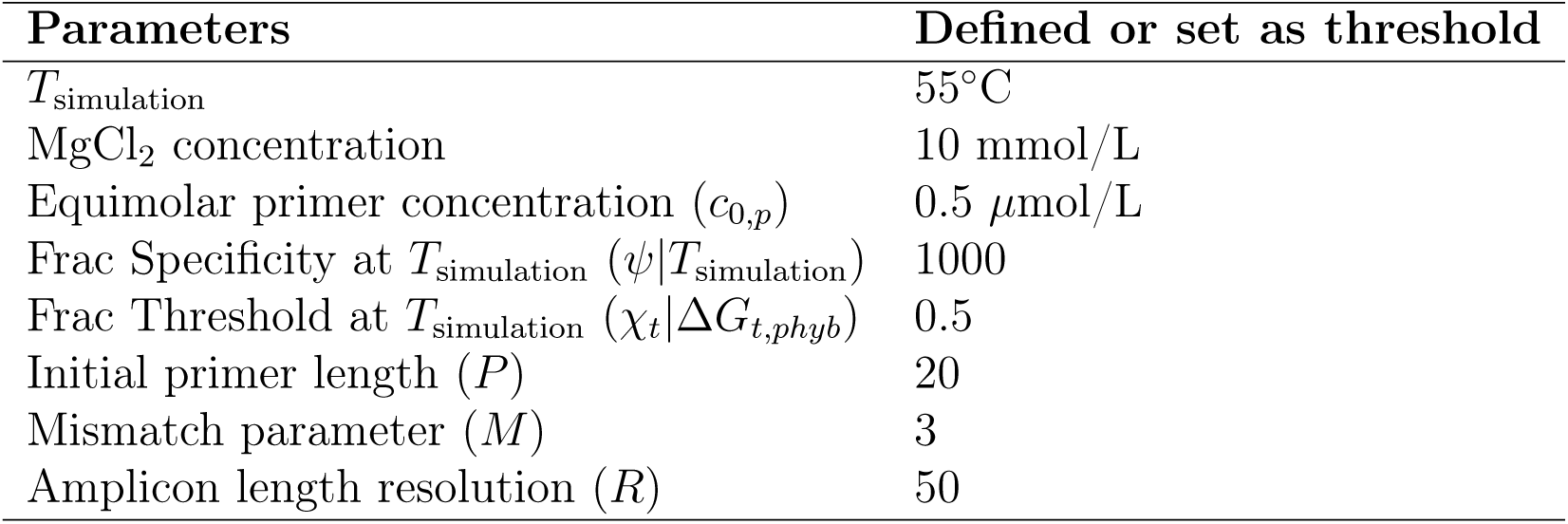
ThermoPlex parameters used in the study for the design of Multiplex PCR Primers.

### 2.4 Laboratory Validation

Performance of the predictive capabilities of the *SimulEq*algorithm in simulating multireaction equilibrium primer interactions was tested using an in-house multiplex assay for identifying five (5) sardine species through the amplification of Cytochrome C Oxidase I (COI) gene. A simulation was performed using the primer sequences listed in Table 2 and then compared with gel electrophoresis profile of the Assay. Meanwhile, the capabilities of the *ThemoPlex* algorithm to assist in the screening of target-specific multiplex PCR primers were also assessed. Multiplex PCR primers (Table 2), also targeting the COI gene region, were designed for an assay to delineate four (4) sardine species using the algorithm. One of the three predicted set of primers were then randomly picked to be evaluated in the laboratory for its performance.

**Table 2:**
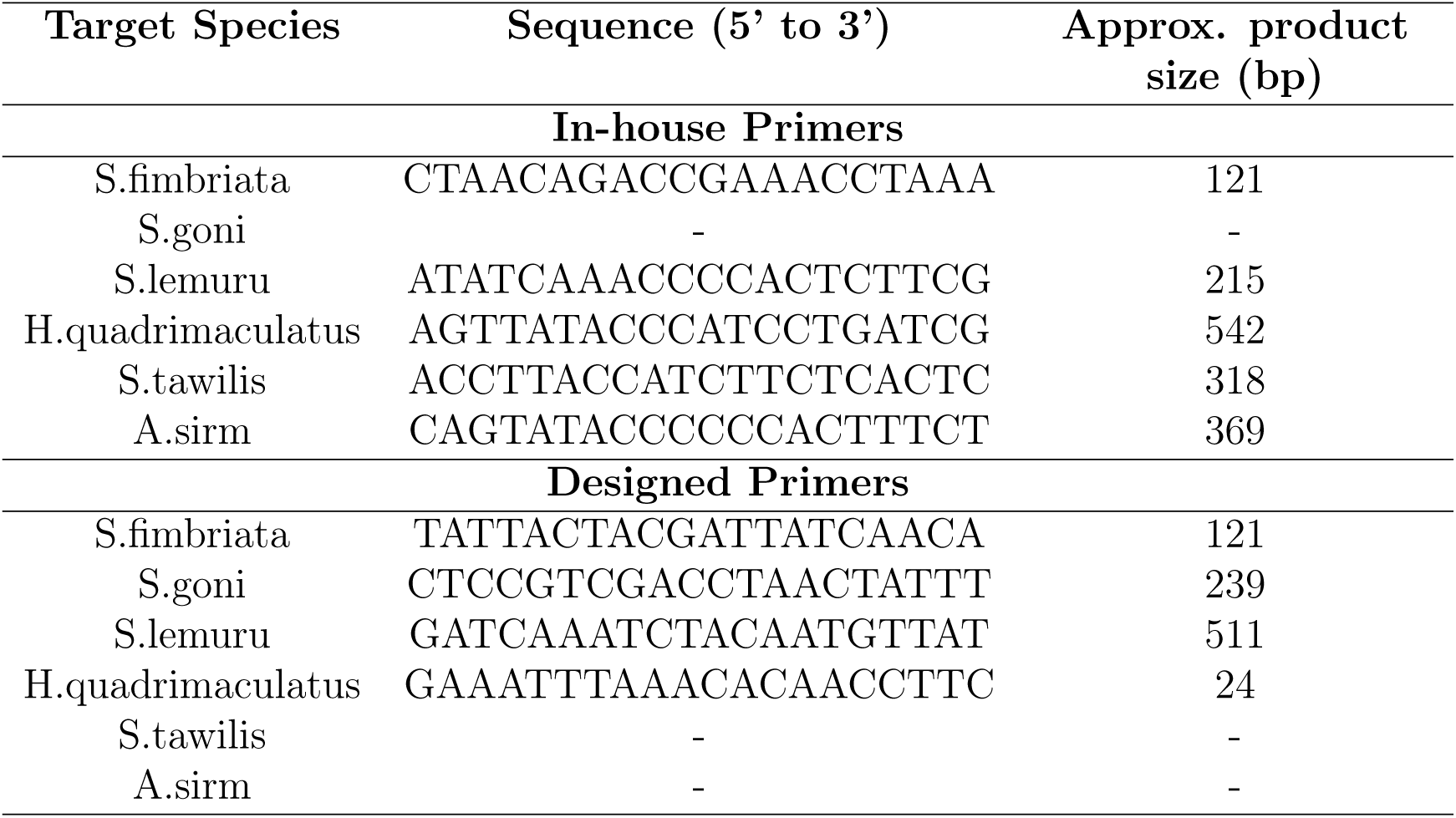
Target species sample set for the PCR trial run with corresponding specific primers and expected amplicon product size for both In-house and ThermoPlex-designed primers.

High-quality DNA used for the experiment were extracted from tissue samples of each of the target species using Qiagen’s DNeasy Blood and Tissue Kit. The extracted DNA was amplified through PCR using a 10 *µ*L PCR mix comprised of 2.45 *µ*L of ultrapure water, 2 *µ*L of 5x GoTaq® PCR Buffer, 0.4 *µ*L of 25mM MgCl_2_, 0.8 *µ*L of 2.5mM dNTP, 5 x 0.5 *µ*L of 10 µM each of the five forward primers, 0.5 *µ*L of 10 *µ*M reverse primer, 0.8 *µ*L of Bovine Serum Albumin (BSA), 0.05 *µ*L of 5 u/*µ*L GoTaq® polymerase and 0.5 *µ*L of DNA template. Gel electrophoresis was then performed using 3% agarose gel to view the results of the amplification.

## 3 Results and Discussion

### 3.1 Benchmarking of *ThermoDHyb* algorithm and *ThermoPlex* subroutines

Table 3 provides the list of model equations used to assess the computational complexity of the *ThermoDHyb* algorithm, and their corresponding fitted parameters. Visual inspection of the results of the time-complexity benchmark (Figure 6) suggests that the quadratic model is the best fit for the resulting simulation runtimes as it demonstrates good overlap with the data points relative to other polynomial models. Moreover, the *b* fitting parameter (*y*-intercept) of the quadratic model has the closest value to zero suggesting the model’s ideal fit compared to others. All these indicates an average-case quadratic time complexity (*O*(*N* ^2^)) for the ThermoDHyb algorithm, for the special case of equal sequence length (*M* = *N* ). The general case is thus expected to be *O*(*M* × *N* ) for *M* ≠ *N* as this represents the computational bottleneck of filling up the *m* × *n* matrix of doublet interactions. The sub-routine’s efficiency better fits the iterative nature of the ThermoPlex algorithm compared to other published algorithms predicting nucleic acid thermodynamic interactions such as McCaskill’s [30] and Dirks’s [31] both running on *O*(*N* ^4^) time. Although their algorithm’s sampling of possible microstates in the ensemble is more rigorous, the microstates predicted in *ThermoDHyb* should be enough to sample the most important contributors in the ensemble free energy, particularly in the case of PCR applications where interactions are limited to small oligonucleotide primers that seldom adopt more complex secondary structures and interactions such as pseudoknots. Indeed, the benchmark results (Table 4) manifest the algorithm’s efficiency, with subroutines completing the computations under 10 minutes across various personal computers and operating systems tested in the study.

**Figure 6:**
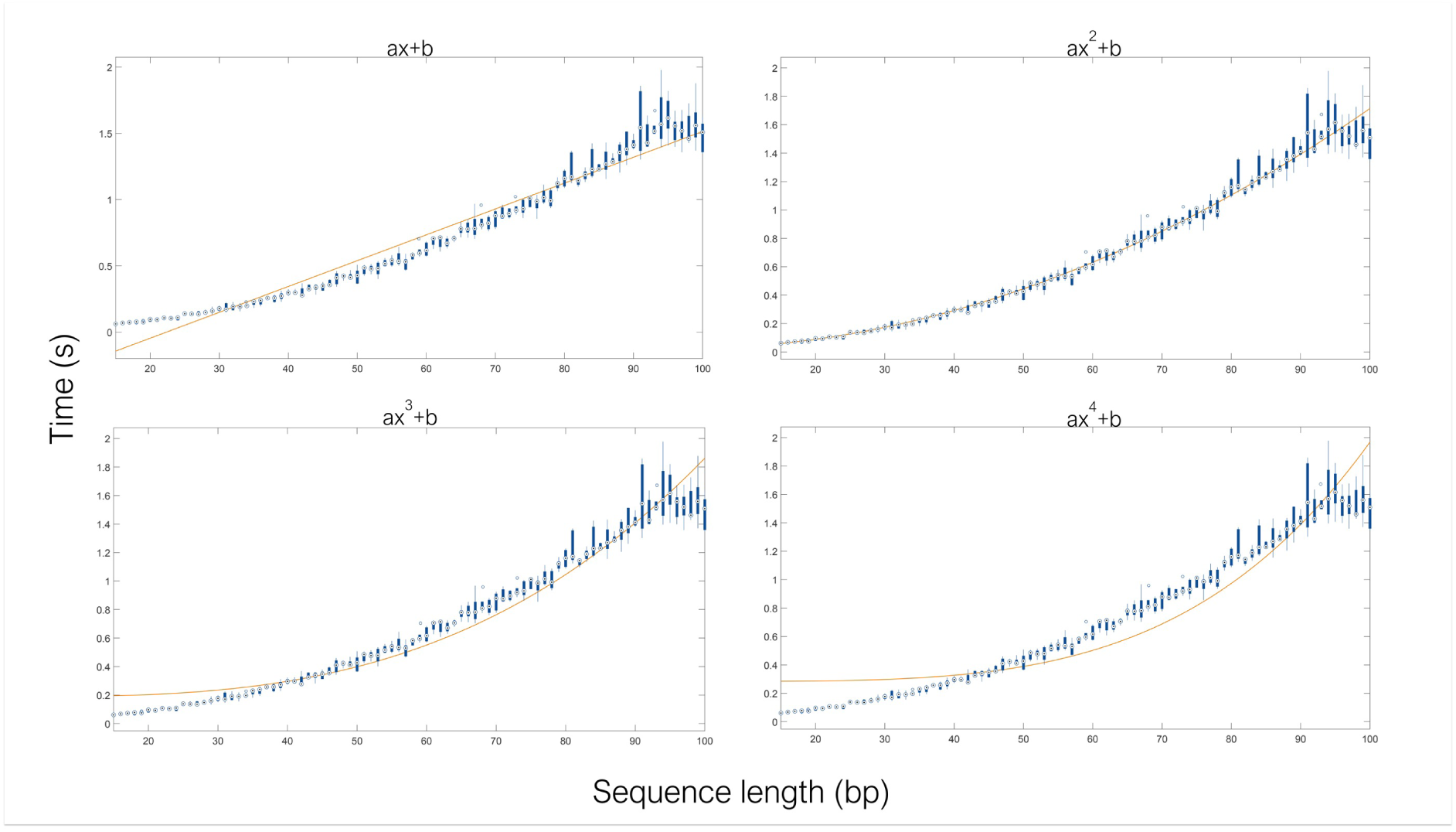
Benchmarking for time-complexity of *ThermoDHyb* through exploration of runtime as a function of sequence length (bp). Box plots (blue) were generated from sequence length of 15 to 100 bp at 1 bp increment with 5 replicates each bp. The orange lines represent each polynomial model fitted through non-linear least squares.

**Table 3:**
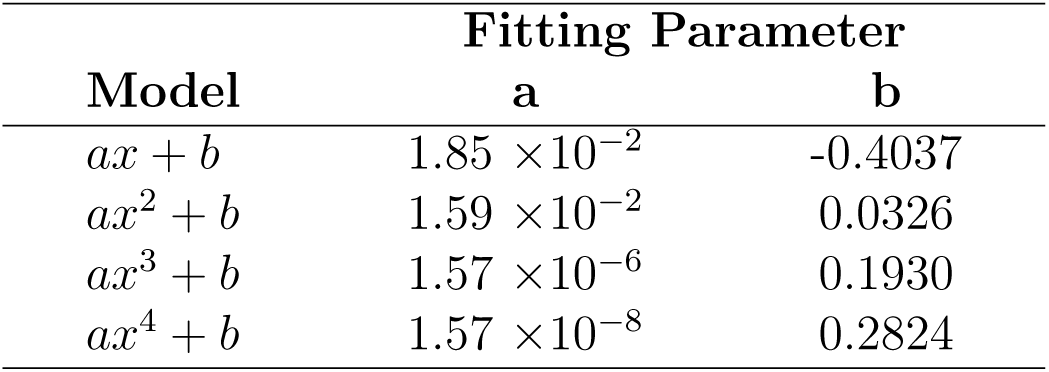
Resulting fitting parameters for each model through non-linear regression of ThermoDHyb runtimes.

| Model | Fitting Parameter |  |
| --- | --- | --- |
|  | a | b |
| $ax + b$ | $1.85 \times 10^{-2}$ | -0.4037 |
| $ax^2 + b$ | $1.59 \times 10^{-2}$ | 0.0326 |
| $ax^3 + b$ | $1.57 \times 10^{-6}$ | 0.1930 |
| $ax^4 + b$ | $1.57 \times 10^{-8}$ | 0.2824 |

**Table 4:**
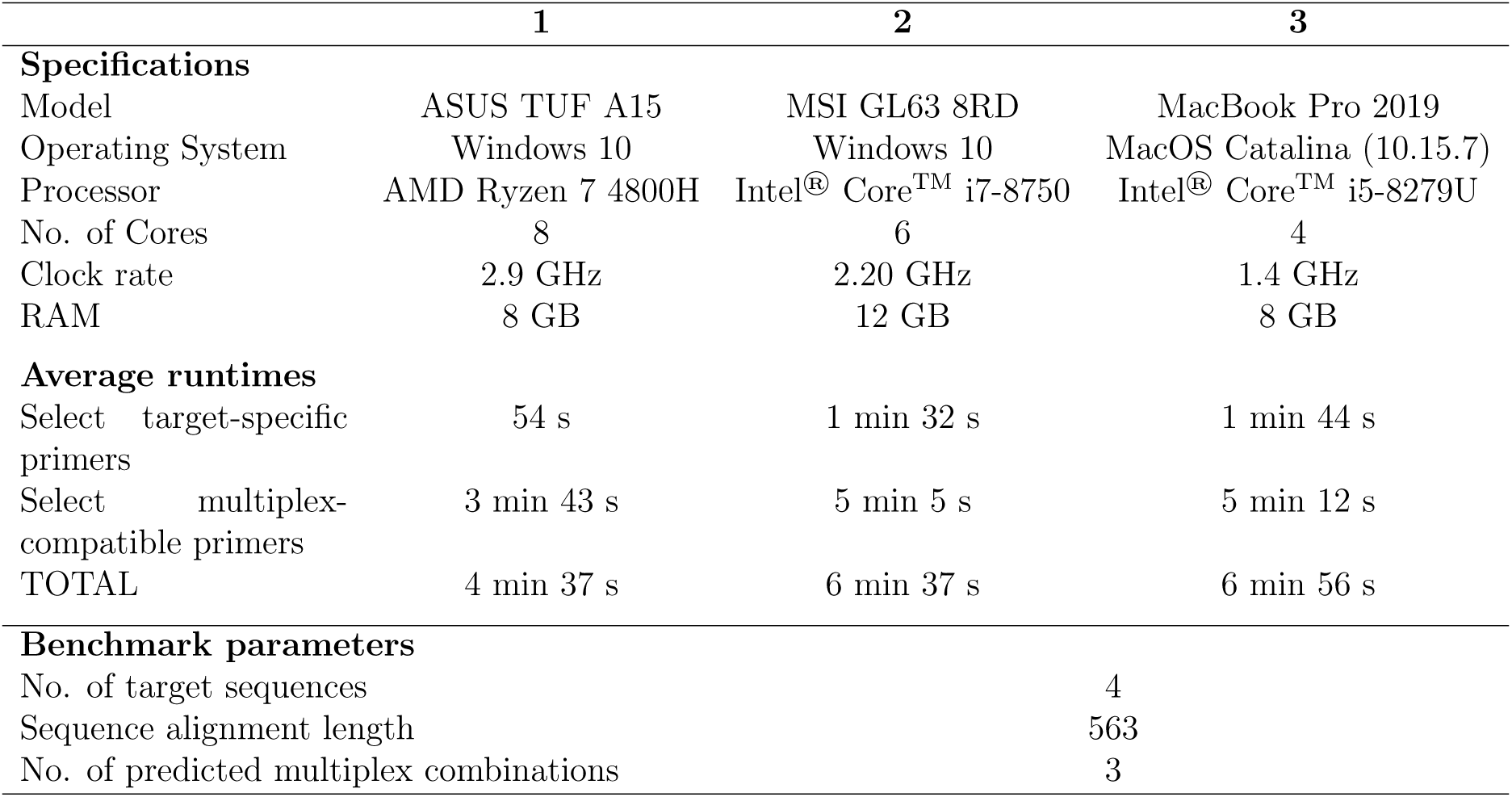
Computing platform specifications, subroutine average runtimes (3 replicates), and input parameters for the ThermoPlex benchmarking.

Meanwhile, results of the linear regression analysis (Figure 7) between ThermoDHyb and three other software (Visual OMP [32], (B) DINAMelt [33], and (C) NUPACK [34]) suggest good concordance in the predicted Δ*G* values, especially with Visual OMP. A near-unity regression slope and *R*^2^ of 0.9959 and 0.9770, respectively, as well as a near-zero *y*-intercept of 0.0192 indicates a good one-to-one agreement in the predicted values. Good concordance (84%) in terms of predicted secondary structures is also observed, with non-concordance existing only at higher Δ*G* values (*>* −3 kcal/mol). These suggest that the *ThermoDHyb* algorithm is at least at par with the accuracy of the Visual OMP software in terms of predicting the thermodynamic interactions of two oligonucleotides. Although quite expensive, the Visual OMP software has been extensively used in numerous multiplex-PCR primer design studies [35, 36, 37, 38] and yielded accurate results. Moreover, fairly good concordance in predicted secondary structures (70% and 84%) and Δ*G* values, at least in terms of trend, can still be seen in the comparison with both DINAMelt and NUPACK. It is also worth noting that the y-intercept of Figure 7C implies a shift in the predicted value of 3.41 kcal/mol on average by NUPACK. This shift is consistent with the study of Zhang et al. [39] where they needed to adjust the NUPACK-predicted Δ*G* values by 1.5 kcal/mol for their model to better fit the experimental results. All of these results establish the reliability of the ThermoDHyb algorithm and thus ensuring a solid framework for thermodynamic calculations of oligonucleotide interactions performed within the ThermoPlex routines.

**Figure 7:**
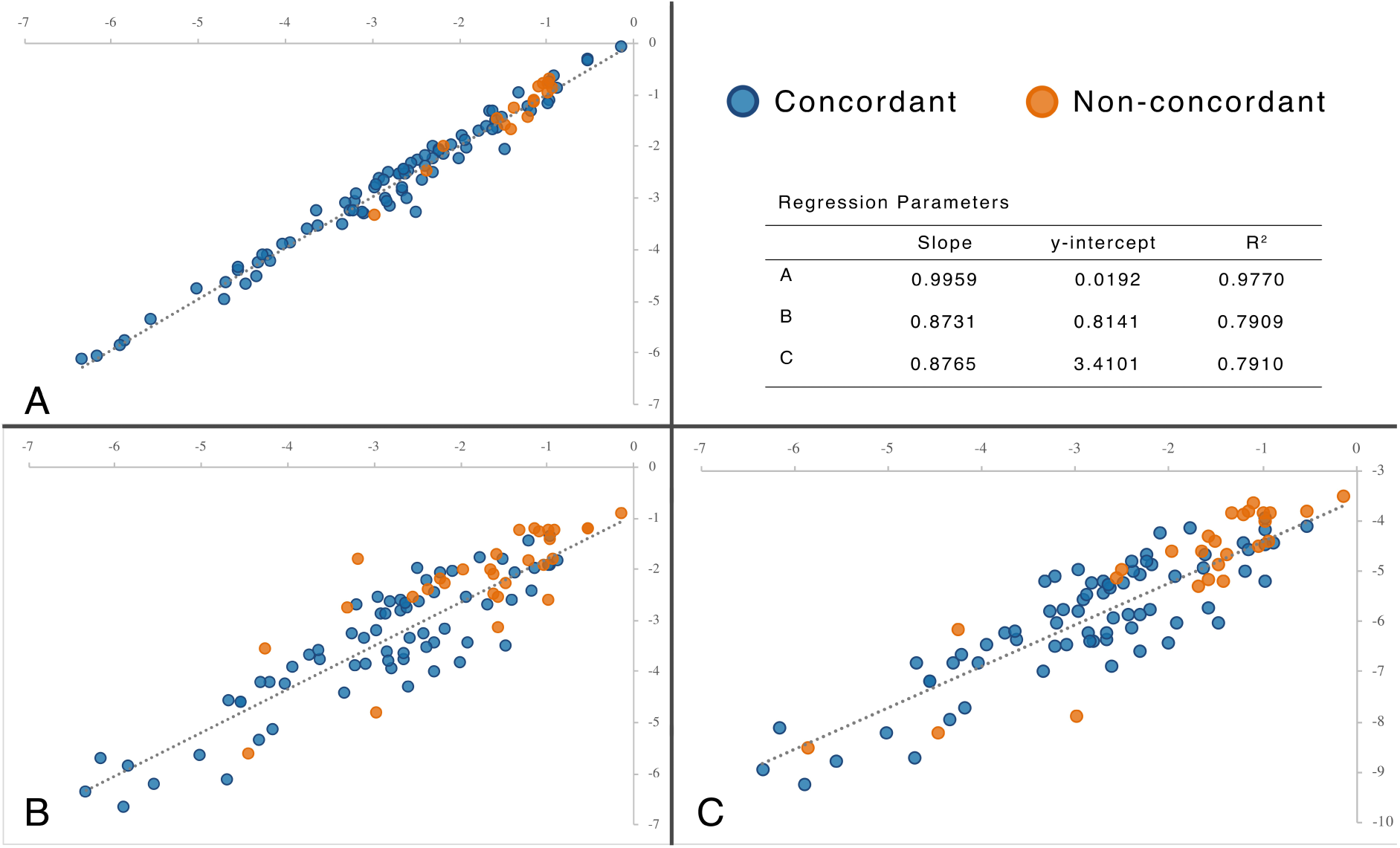
Linear regression analysis for the benchmarking of predicted global minimum free energy values (kcal/mol) simulated at 37^◦^C and 1M NaCl solution as predicted by *ThermoDHyb* as compared to various software: (A) Visual OMP [32], (B) DINAMelt [33], and (C) NUPACK [34]. Blue and orange data points correspond to concordant and non-concordant predicted secondary structures between *ThermoDHyb* and the software.

### 3.2 Sensitivity analysis of the ***χ*** model for fractional conversion

The scatter plots from Monte Carlo (MC) simulations in Figure 8A-C show that *χ* is most sensitive to temperature parameter changes, followed by MgCl_2_ concentration, and lastly *α*. This is elucidated by a more distinct pattern in the plot of C than in B, and then in A. Such is an expected trend since DNA hybridization is most strongly influenced by temperature as heat is mainly responsible for breaking the hydrogen bonds of the DNA’s N-bases in PCR reactions. Meanwhile, salt concentration moderately influences DNA hybridization stability through the neutralization of the negative charges on the backbones by cationic metals (Na^+^ and Mg^2+^), reducing electrostatic repulsion between phosphates [40]. On the other hand, no pattern is discernable in plot A, but further evaluation was performed since the pattern may be heavily diluted by high variations in *T* . This was done by performing another MC simulation where *T* is held constant at 62°C (Figure 8 D-E), thus comparing only the influence of MgCl_2_ and *α* parameters. The results of the second simulation depict that although a pattern is completely discernable in plot E and still none in plot D, outliers (yellow) can be observed in the former indicating that *α* still influences the model to a certain extent. However, linking these outliers to plot D (also yellow) suggests that the significant influence of *α* is limited only to lower limit values (*α <* 30), while the influence diminishes as values of *α* goes higher (blue). PCR reactions are typically conducted with *α* of at least 108 to ensure that template will anneal to primers rather than itself [41], thus the model is sufficiently valid for PCR applications despite the uncertainties attributed to *α*. The simulation results also provide an interesting insight in the context of modelling any bimolecular interactions beyond PCR applications, as the *χ* function was derived from a generalized mass action equation for bimolecular reactions. The model can thus be used for predicting fractional conversion of a limiting reactant even without accurate information about its value, given that the excess reactant is at least at a factor of 102 more than the limiting.

**Figure 8:**
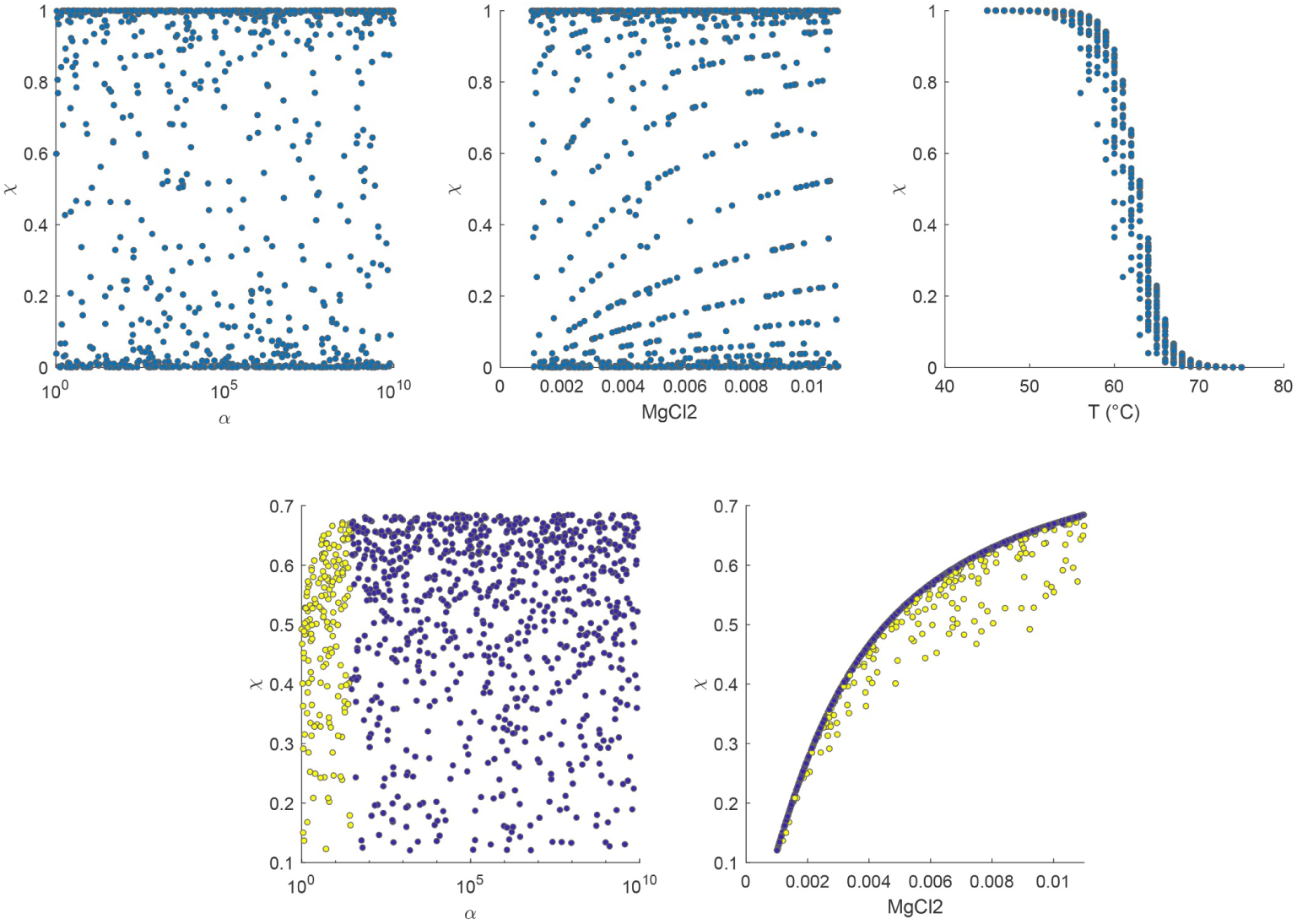
Monte Carlo (MC) simulation (*n* = 1000) scatter plots of *χ* versus *α* (A and D), MgCl_2_ concentration (B and E), and T (C). The parameter *T* was held constant at 62^◦^C for MC simulations (*n* = 1000) in D and E.

### 3.3 *SimulEq* and *ThermoPlex* Simulations, and Laboratory Validation

Generated EPD curves from Figure 9 through the *SimulEq* algorithm illustrates the predicted product distributions of the whole multiplex assay as functions of temperature. Each plot corresponds to the multiplex assay’s performance on each target species’ gene sequence. Inspection of the resulting EPD curves suggest that the assay’s primers for *S. lemuru* and *S. fimbriata* are completely specific to their own respective target species. This is because in plots D and E, apart from products from their respective targets, there are no other curves visible. Such is the opposite to plots A to C as multiple curves are visible aside from each of their own target curves, particularly the primer for *S. tawilis* (plot B) wherein multiple curves are already obvious for as a high a temperature as 55^◦^C. This implies that the primer for *S. tawilis* should demonstrate a certain degree of non-specificity towards *S.lemuru* and *S. fimbriata*. Indeed, this is consistent with experimental results as shown in Figure 11A. Gel electrophoresis profile for the assay indicates the same expected non-specificity of *S. tawilis* primers for both species, as shown by the two faint gel bands in its lane whose sizes are consistent with that of *S. lemuru* and *S. fimbriata* PCR products. The consistency of *SimulEq* simulations with experimental results demonstrates the reliability of the algorithm in the thermodynamic analysis of multiplex PCR assays. It is worth noting, however, that this type of theoretical analysis does not directly translate to expected PCR products per se due to the simulation capturing only what is happening per annealing cycle, or the first cycle at best. Nevertheless, the analysis done on the EPD curves still proves to be insightful since it illustrates the assay’s behavior when hybridization due to non-specific primer binding becomes more and more significant at lower temperatures.

**Figure 9:**
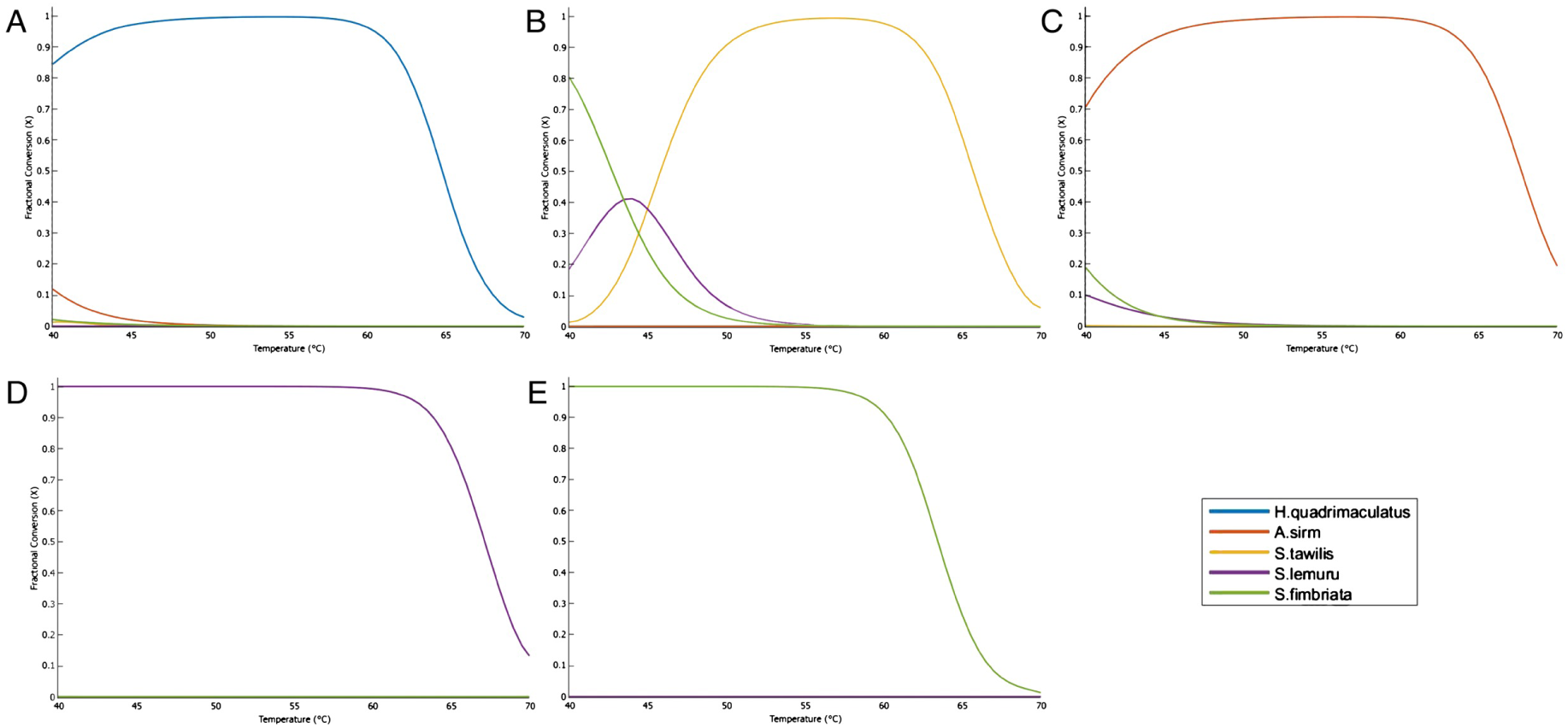
Equilibrium Product Distribution (EPD) curves as function of temperature generated by *SimulEq* from the in-house developed multiplex PCR assay primers simulated per target COI gene sequence. Each color corresponds to the target species the primer is designed for. Primer sequences and their targets are listed in Table 2.

The output of the ThermoPlex design routine is shown in Figures 10 and 11 which includes the multiplex specificity (*ψ_i_*(*T* )) curves and the predicted gel electrophoresis profile for the designed assay. Due to the unavailability of a scoring function (still under development) to rank all the designed candidate multiplex primer sets, random selection of one assay out of three candidate primer sets were done for laboratory validation. Following the same analysis of the EPD curves as above suggests that the primers in the designed assay are expected to be more target-specific than the in-house developed primers. The *ψ_i_* curves in Figure 11B also aids in this inference as it illustrates the expected fractional specificity of the primer set at various temperatures. It shows the temperature limits at which each primer in the assay should theoretically retain its specificity above the user-defined *ψ* threshold (black horizontal line). Moreover, the curves suggest that at Tsimulation of 55^◦^C, the assay retains specificity, with the least specific primer (*S. lemuru*) still having *ψ_i_* values in the order of 104, i.e., the primer is expected to hybridize 104 times more to the target than all the other primers in the assay combined. The result of the simulation is indeed supported by the experimental results in Figure 11B. The gel electrophoresis profile of the designed multiplex assay shows no discernable non-specificity, with only one distinct gel band manifesting per each target. The experimental results thus show that the set of multiplex primers automatically designed by the *ThermoPlex* routine performs adequately, proving the reliability of the primer screening routine the software implements. Tests involving larger data sets (e.g. longer sequence length, more targets), however, should be performed to further assess the automated screening capabilities of *ThermoPlex*.

**Figure 10:**
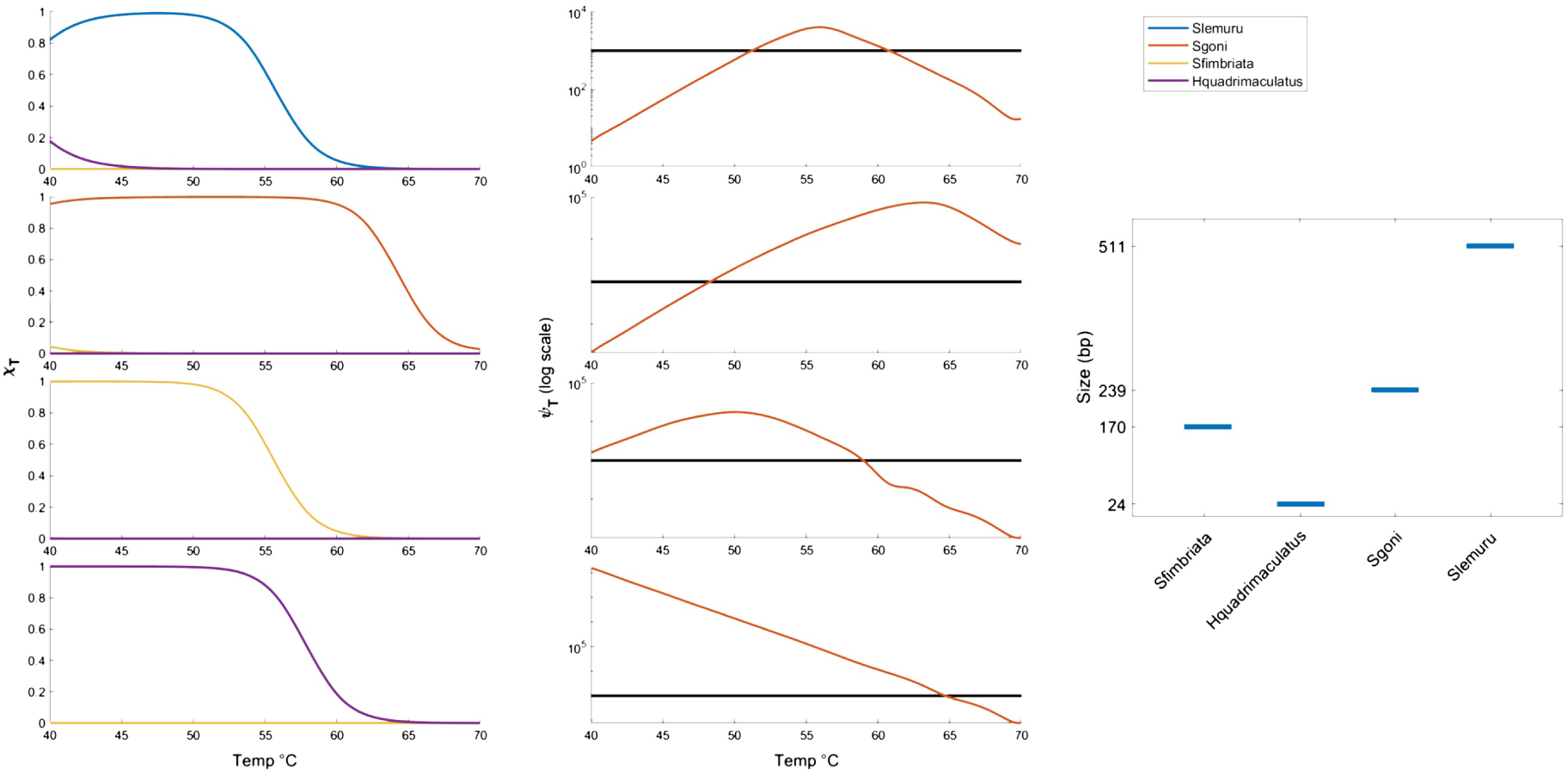
Output of the *ThermoPlex* ’s design algorithm showing A) EPD curves, B) *ψ*(*T* ) curves (log10 *y*-axis scale) as functions of temperature (black lines correspond to *ψ* = 1000), and C) predicted gel electrophoresis profiles for the multiplex assay.

**Figure 11:**
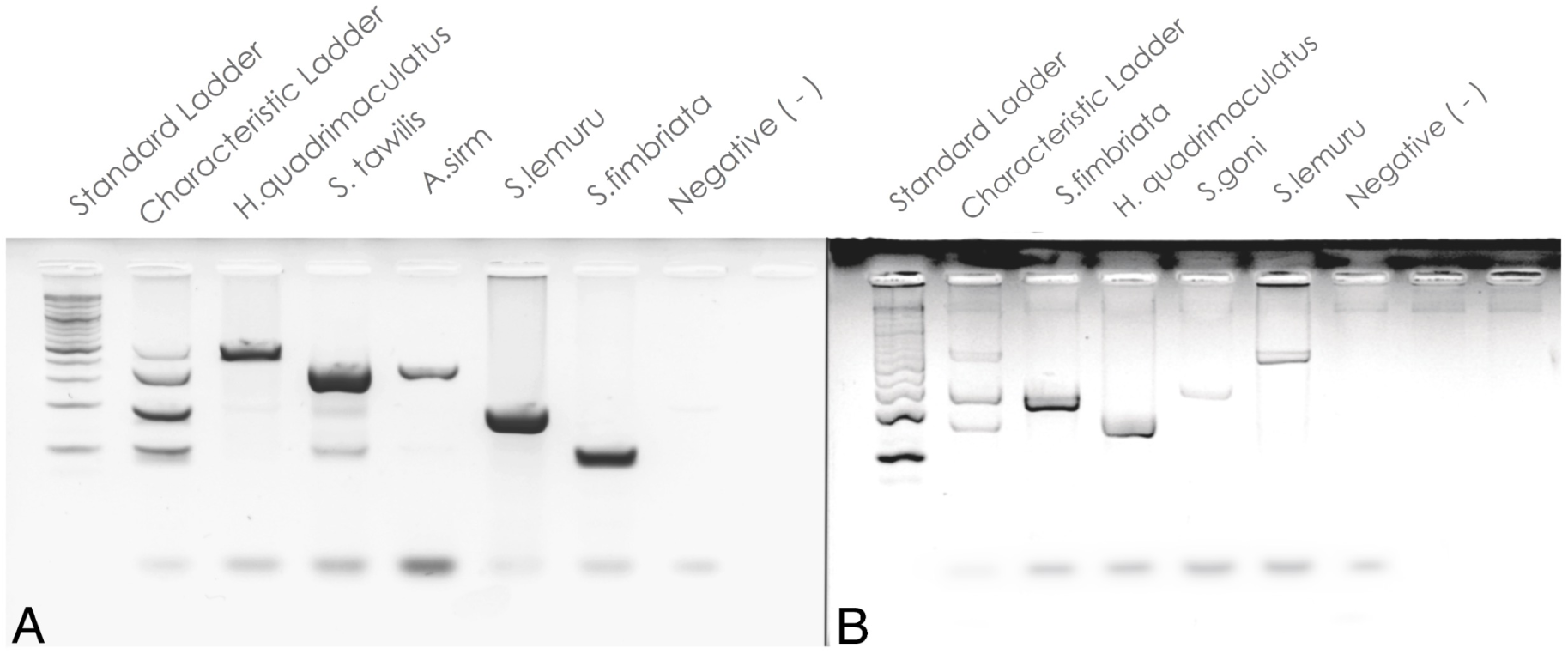
Gel electrophoresis profile of the PCR products for A) in-house developed multiplex PCR assay and B) *ThermoPlex* -designed the multiplex PCR assay.

## 4 Conclusion and recommendations

Despite the proved performance of ThermoPlex in automatically designing multiplex PCR primers, one caveat is the limitation in terms of sequence input to the program, specifically:

1. The program strictly needs the sequence input to be aligned and trimmed to equal length.
2. In its current state, it is unable to handle sequence alignments with gaps, i.e., gene regions with insertions and deletions.
3. It also fails to pick candidates for sequences with highly dispersed and sparse polymorphic sites despite having fairly high genetic distances.

Moreover, the laboratory performance assessments so far of ThermoPlex and all of its subroutine lack sufficient quantitative basis despite the program being motivated by quantitative thermodynamic variables. Hence, it would be best to test its predictions with quantitative experiments such as qPCR-based methodologies. These types of experiments would also empirically fine-tune the models underlying the algorithm’s framework, making them more consistent to real-life expectations.

Input-parameter optimization was also not done in order to exhaustively explore the output candidate primer space to design more optimal multiplex primers. Additionally, a ranking/scoring system would also be needed, especially if the program were able to select numerous candidate primer sets that will be difficult for the user to evaluate and handle. Presently, the workaround would be to make the parameters (e.g., MgCl_2_ concentration, specificity thresholds) stringent enough to produce a manageable number of candidate sets. All of these will be implemented in the future to further optimize the selection process.

In summary, the ThermoPlex program demonstrates great potential to improve the accuracy and speed of designing multiplex PCR primers through all the evidence presented in this study. With the program’s automated screening routine, designers can expect to have candidate multiplex PCR primers within minutes, subject to the speed of the computing platform, number of design targets, length of the input sequence, and number of candidate multiplex set predicted. All these are achievable through common personal computers without the need for high performance computing. Moreover, apart from the automated primer screening algorithm, the program also offers useful and insightful tools motivated by DNA thermodynamics via its ThermoDHyb and SimulEq routines. Depending on the needs of the user, these novel algorithms can be used independently or in concert with other existing programs. Although already useful in its current state, further improvements in the future are to be implemented to enhance the program’s capabilities such as more robust handling of sequence alignment, ability to score and rank predicted multiplex sets, and possible expansion to a web server format.

## 5 Availability

The standalone app together with the source codes are available in the GitHub repository (https://github.com/aagmata/ThermoPlex)

## 6 Acknowledgement

The authors would like to acknowledge the contribution of Dr. Ma. Rio. Naguit, Dr. Asuncion De Guzman, Mr. Jerry Garcia, Mr. Jhunrey Follante, and Mr. John Christopher Azcarraga in the collection and processing of sardine samples. Additionally, the authors would like to express their gratitude to Dr. Dame Loveliness Apaga-Agmata for her valuable insights in chemical thermodynamics and physical chemistry.

