## Supplementary Material for "ThermoPlex: An Automated Design Tool for Target-specific Multiplex PCR Primers based on DNA Thermodynamics"

In this supplementary material, we will be discussing in detail the algorithms used in the study, and the assumptions, equations, derivations, and models that serve as the foundation for the computational methods presented.

**Nomenclature**

| | |
|---|---|
| **Variables** | |
| $T$ | Temperature (K) |
| $C_P$ | Heat capacity at constant pressure |
| $\Delta G$ | Standard Gibbs free energy |
| $\Delta H$ | Standard Enthalpy |
| $\Delta S$ | Standard Entropy |
| $n$ | Number of moles |
| $Z$ | Partition function |
| $c$ | Molar concentration (mol/L) |
| $\lambda$ | Lagrange undetermined multiplier |
| $\chi$ | Fractional conversion |
| $\psi$ | Fractional specificity |
| $\alpha$ | Primer-template concentration ratio |
| $P, P'$ | Cumulative pairwise energy matrix |
| **Indices** | |
| $i, j$ | Single-stranded species index |
| $k$ | Double-stranded species index |
| $o$ | Variable species index in the partial derivative |
| $p$ | Constant species index in the partial derivative |
| $m, n$ | Pairwise doublet index in matrix $P$ |

*ThermoPlex* models the complex interaction of DNA primers in a PCR solution in equilibrium. The program incorporates two algorithms (*ThermoD-Hyb* and *SimulEq*) in selecting target-specific, multiplex-compatible primers. The models used operate according to the following assumptions and limitations:

1. The DNA template in the solution is the limiting reactant in the simulation.

2. The models are established around the first annealing event of the PCR reaction cycle.

3. Primers binding along the extension path of another primer completely inhibit the amplification product of the latter.

### 1 ***ThermoDHyb*** Algorithm

The *ThermoDHyb* algorithm aims to predict the thermodynamics of the double-stranded interaction between two single-stranded DNA, and the associated Gibbs free energy change ($\Delta G$) from a single-stranded reference state towards each optimal/suboptimal microstates. Among these microstates the corresponding double-stranded secondary structures of each optimal state (minimum free energy state) is also predicted. The algorithm derives its working principle from the Smith-Waterman algorithm [1] for sequence alignment. Sequences $i$ and $j$ with lengths $M-1$ and $N-1$, respectively, are compared using an $M$ by $N$ pairwise energy matrix $P$ (Figure 1) of cumulative $\Delta G$ per sequence doublet according to the Nearest-Neighbor model parameters provided by SantaLucia et al. [2] The values in the table are filled iteratively according to this equation:

$$\Delta G_{m,n} = \min \begin{cases} \Delta G_{bu(m,n)} + \Delta G_{m-bu,n-1} + \Delta G_{\text{bulge}(1:bu)}, \\ \Delta G_{bl(m,n)} + \Delta G_{m-1,n-bl} + \Delta G_{\text{bulge}(1:bl)}, \\ \Delta G_{\text{NN}(m,n)} + \Delta G_{m-1,n-1} + \Delta G_{\text{loop}}, \\ 0 \end{cases} \tag{1}$$

The iteration starts at the $m$th row and $n$th column $(m, n)$ of $P$ and requires the $(m-2, n-1)$, $(m-1, n-2)$, and $(m-1, n-1)$ elements to be filled up initially. The dependency ends at $(1, m)$ or $(n, 1)$ where terminal energy calculations according to the Nearest-Neighbor (NN) model will be performed. The value of $\Delta G_{m,n}$ is determined by equation 1 depending on the case that will give the minimum value. The evaluation in each iteration is done against logical statements corresponding to rules governing DNA structural motifs and their energies (e.g. internal loop formation and propagation, bulges, etc.)[2].

Furthermore, the algorithm generates two of such matrix $P$ and $P'$, one for each direction of propagation (Figure S2). This was done to properly incorporate terminal interaction energies in both ends of the DNA structure, as the procedure recognizes only the initiating end of the structure along the propagation path. Consolidating the initiating doublet energies on both propagation direction should arrive on an equivalent total interaction $\Delta G$ since the Gibbs energy parameter is a state variable/function and is hence independent of the path taken between initial and final states. The consolidation involves a recursive path-tracing algorithm that starts at an address in P that is the same as the initiation matrix coordinates in $P'$ (as depicted by the orange box and arrows in Figure 2). The program then traces its way (green arrows) towards the initiation coordinates in P, then starts again at the same coordinates in $P'$ tracing back towards the initiation coordinates

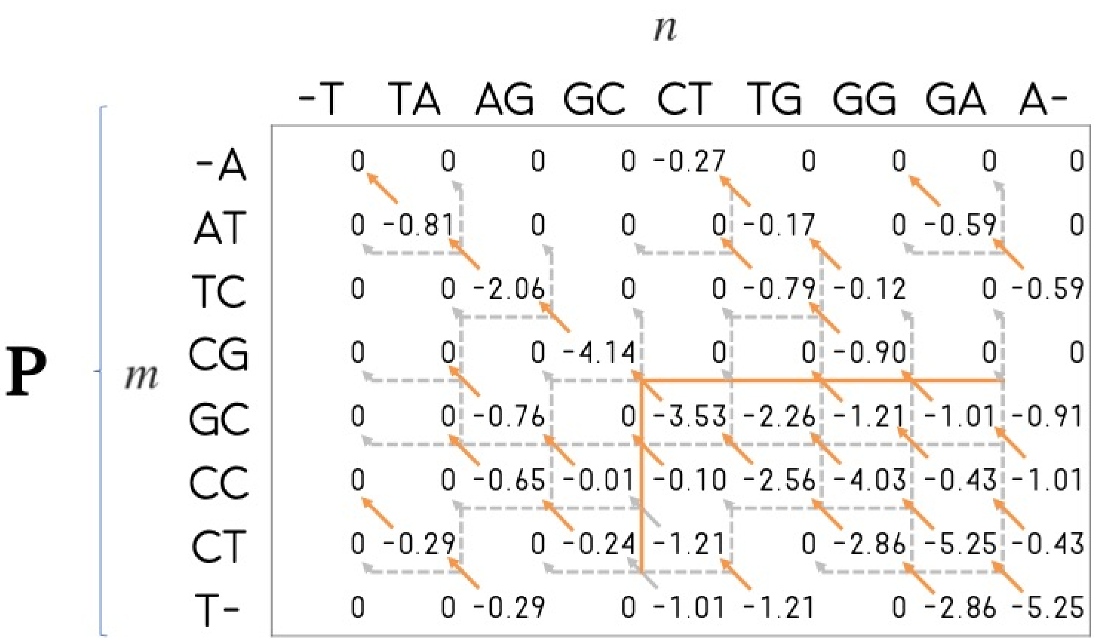


**Figure 1:** Pairwise energy matrix of doublet pairs for sequences 5' ATCGCCT 3' (Sequence i) and 5' AGGTCGAT 3' (Sequence $j$) at 40 °C and 1M NaCl simulation conditions. Sequence $i$ and $j$ doublets are listed per row and column respectively. Arrows represent the cumulative dependence of each element in $(m, n)$ to $(m - bulge, n - 1)$, $(m - 1, n - bulge)$, or $(m - 1, n - 1)$ elements. Orange, solid arrows depict the dependence case chosen.

in $P'$. This forms a loop when the start and end addresses in $P$ and $P'$ are the same, a criterion which the program looks at when seeking multiple true local energy minima.

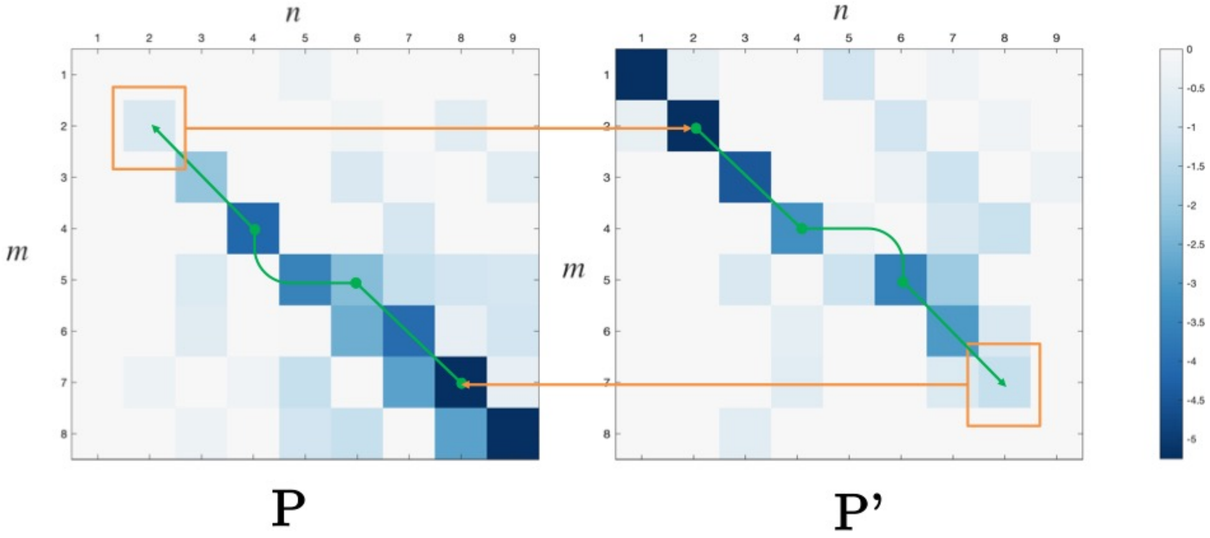


**Figure 2:** Propagation of cumulative energy scores (down-right for $P$, up-left for $P'$). Terminal doublet pairs are boxed in orange, green arrows show the path of tracing the doublets for inferring the minimum free energy (MFE) structure.

The resulting local minima of cummulative scores per propagation path are stored and

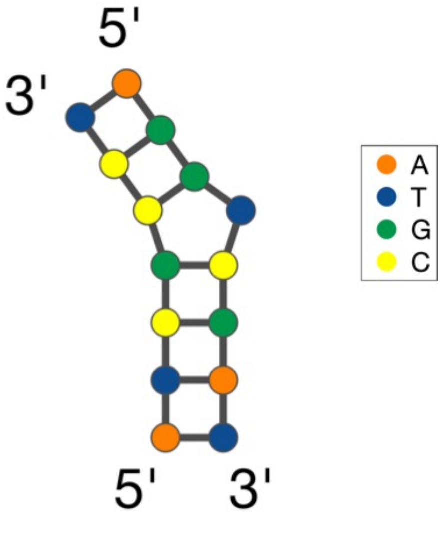


**Figure 3:** MFE structure and corresponding free energy change $\Delta G^{i,j}_{\text{hyb}}$ predicted by *ThermoDHyb* for sequences 5' ATCGCCT 3' and 5' AGGTCGAT 3' at 40°C and 1M NaCl simulation conditions.

used for the calculation of the system partition function $Z$. Defined in equation 2, the system partition function is the sum of the statistical weights for all the possible secondary structures - defined as the system microstates - that can exist through the interaction between two strands [3]. Here, we assumed that the only complex microstate (system and complex microstate definition according to Schaeffer et al.[4]) that exists for each system microstate is the local minimum free energy state resulting from a two-state approximation of each system microstate. Such approximation is proven to be accurate enough to predict thermodynamic properties of short oligomers [2]. In contrast though with models purely relying on the two-state approximation assumption, our model includes the existence of various system microstates contributing to the system Gibbs energy. This contribution is accounted for by deriving the interaction Gibbs energy value from the partition function using equation 3.

$$Z_{i,j} = \sum_{s \in \sigma} e^{\frac{\Delta G^{i,j}_s}{RT}} \tag{2}$$

$$\Delta G^{i,j}_{\text{hyb}} = RT \ln(Z_{i,j}) \tag{3}$$

However, as a limitation of the *ThermoDHyb* algorithm, the local microstates predicted are limited only to optimal structures corresponding to different random nucleation sites. The algorithm fails to predict distinct states existing on the same nucleation site, a consequence of trying to abate the computational complexity of the calculation. In such cases, the algorithm will choose the path leading to the most optimal structure per nucleation site. However, we believe that the state optima sampled by the algorithm through the interaction matrix is enough to sample all the significant contributors to the partition function. The advantage of this algorithm over pre-existing algorithms such as McCaskill's [5] algorithm for recursive computation of the interaction partition function is that the algorithm runs only on $O(M \times N)$ time complexity (quadratic for $M = N$), suitable for the iterative nature of the downstream procedures to be described in this study. The algorithm also does not account in its thermodynamic calculations the secondary structures that arise from intra-strand interaction as this happen less frequently on short oligonucleotides than in longer ones, with the former being used commonly as primers for PCR-based applications. But since it is still an important parameter for selection of PCR primers in general, it is incorporated in the selection process as a heuristic before thermodynamic calculations are performed. This is performed through MATLAB's bioinformatics toolbox function *rnafold*. Presence of predicted secondary structure through this function, regardless of thermodynamic stability, will result to rejection of the primer candidate.

### 2 *SiMulEq* Algorithm

*SiMulEq* simulates the complex equilibrium scenario of interacting nucleic acid species in a Multiplex PCR reaction tube by modeling the event as a set of parallel competing reactions (4). The same model has been demonstrated by SantaLucia et al. [2] with the goal of determining the effective melting and thermodynamic parameters given that the desired interaction is hindered to a certain extent by all the competing nucleic acid strands present in the reaction mix. They also used it to predict the equilibrium product distribution curves of any species of interest inside the same mixture. In our case, we employed a similar model for SimulEq but with a different computational path accounting for statistical thermodynamic models of chemical potentials [6]. Hence, the problem was converted to that of a total system free energy minimization described by the summability relation of partial properties:

$$nG_T = \sum_i n_i \left[ \frac{\partial (nG_T)}{\partial n_i} \right]_{T,P,n_j} = \sum_i n_i \mu_i \tag{4}$$

The goal is to find a set of $n_i$ values that will minimize the value of $nG_T$. This is because at equilibrium, the system tends towards the minimization of the $nGT$. A constraint can

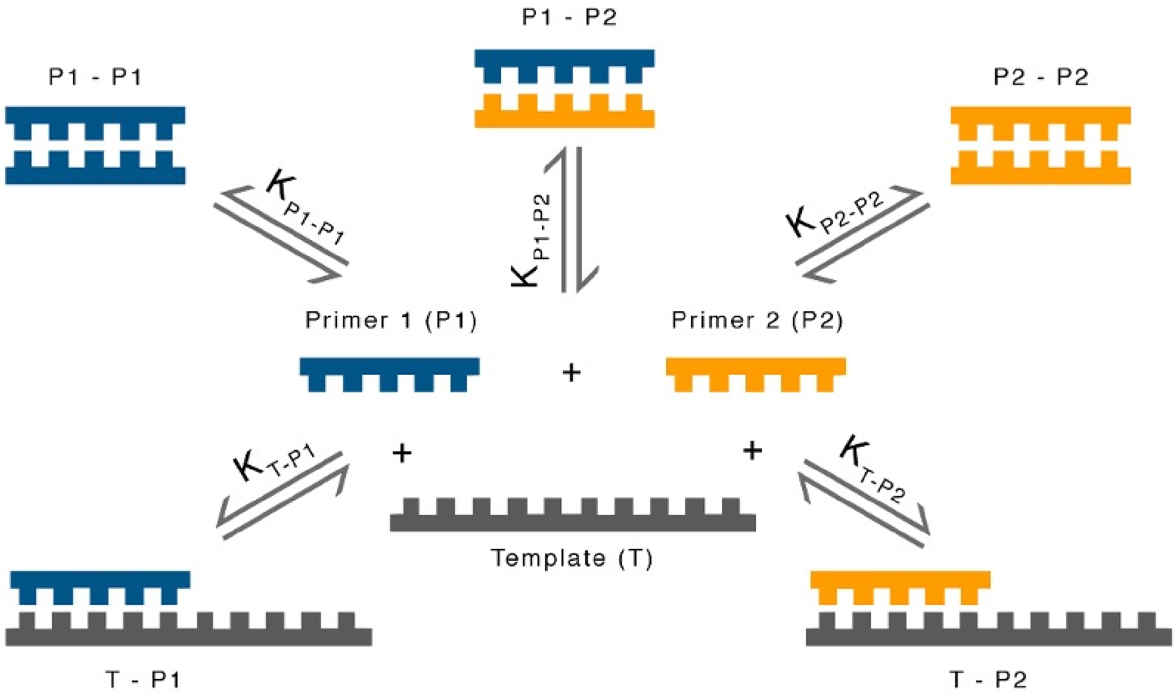


**Figure 4:** Complex multi-reaction equilibrium scenario accounting for each of the possible pairwise interactions in a two-primer, one-template PCR reaction system. Template-template interactions are not accounted for as it occurs on a different mechanism in the PCR annealing step.

be placed on possible solutions to $n_i$ based on the material balance principle, hence the following equation:

$$n_{0,i} - n_i - n_k^{ii} - \sum_k n_k^{ij} = 0 \tag{5}$$

where $n_{0,i}$, $n_i$, and $n_k^{ij}$ correspond to the moles of initial single-stranded, equilibrium single-stranded, and equilibrium double-stranded from single-stranded species $i$ and $j$, respectively. Given a constrained minimization problem, the method of Lagrange undetermined multipliers (LUM) can be employed [7], converting the problem to a multi-dimensional root-finding problem. A Lagrange multiplier $\lambda_i$ is multiplied to the constraint equation corresponding to each of the $i$th material balance equation:

$$\lambda_i \left( n_{0,i} - n_i - n_k^{ii} - \sum_k n_k^{ij} \right) = 0 \tag{6}$$

These equations are summed over $i$, giving:

$$\sum_i \lambda_i \left( n_{0,i} - n_i - n_k^{ii} - \sum_k n_k^{ij} \right) = 0 \tag{7}$$

Addition of this equation to $G_T$ yields a new function $F$:

$$F = G_T + \sum_i \lambda_i \left( n_{0,i} - n_i - n_k^{ii} - \sum_k n_k^{ij} \right) \tag{8}$$

$F$ and $G_T$ have equal values because the term inside the summation is zero. Their partial derivatives, however, are unequal since $F$ incorporates the material balance constraints to $G_T$. The minimum value of $F$ and, equivalently, $G_T$ occurs when all the partial derivatives with respect to $n_i$ are zero, hence:

$$\left[\frac{\partial F}{\partial n_o}\right]_{T,P,n_p} = \left[\frac{\partial G_T}{\partial n_o}\right]_{T,P,n_p} + \sum_i \alpha_i \lambda_i = \mu_o + \sum_i \alpha_i \lambda_i = 0 \tag{9}$$

where index $o$ runs along the totality of double- and single-stranded species $(i + k)$ with $o \neq p$, $\alpha_i = 2$ when differentiating $n_o = n_k^{ii}$, $\alpha_i = 1$ otherwise. Expanding on Dimitrov et al.'s definition [6]:

single-stranded:

$$\mu_i = RT \ln \left( \frac{c_i}{c_{0,i}} \right) \tag{10}$$

double-stranded:

$$\mu_k = \Delta G_{hyb}^{ij} + RT \ln \left( \frac{c_k^{ij}}{c_{0,i} c_{0,j}} \right) \tag{11}$$

Substituting the chemical potential models to the partial differential of F results to the following system of non-linear equations expressed in standard molar concentration units (mol/L):

$$c_{0,i} - c_i - c_k^{ii} - \sum_k c_k^{ij} = 0 \quad (i = 1, 2, \ldots, p) \tag{12a}$$

$$\ln \left( \frac{c_i}{c_{0,i}} \right) + \sum_i \frac{\alpha_i \lambda_i}{RT} = 0 \quad (i = 1, 2, \ldots, p) \tag{12b}$$

$$\frac{\Delta G_{\text{hyb}}^{i,j}}{RT} + \ln \left( \frac{c_k^{ij}}{c_{0,i} c_{0,j}} \right) + \sum_i \frac{\alpha_i \lambda_i}{RT} = 0 \quad (i = 1, 2, \ldots, q) \tag{12c}$$

This is a fully specified system of equations with $p$ material balance equations, $p$ single-stranded and $q$ double-stranded equilibrium equations $(2p+q)$, with $p+q$ of $c_i$ and $c_k$, and $p$ of $\lambda_i$ unknowns $(2p+q)$. Solution to this system is obtained through Newton's method implemented in the *SimulEq* algorithm. *SimulEq* automatically generates the system of equations dynamically according to the given number of initial single-stranded DNA species involved in the reaction mix.

One challenge to using the Newton's method is its poor global convergence. This limitation is very applicable to the problem at hand especially since it is a poorly scaled system. This can be resolved by employing an initial guess that is a good approximation of the true solution for concentration values. This is done using the material balance equations, law of mass action relation and the Gibbs factor. This results to the following system of equations with satisfactory scaling and global convergence with Newton's method:

$$c_{0,i} - c_i - c_k^{ii} - \sum_k c_k^{ij} = 0 \tag{13}$$

$$\frac{c_k^{ij}}{c_i c_j} = e^{-\frac{\Delta G_{\text{hyb}}^{i,j}}{RT}} \tag{14}$$

The resulting concentration values are plugged in as an initial guess to equation system (Equation 12). These are good guesses scaled proportional to the true solution. This allows the Newton's method to be robust in solving the system of equations (Equation 12).

### 3 Fractional Specificity ($\psi$)

The equation for fractional specificity is based on the law of mass action for the reaction corresponding to the DNA hybridization event: template ($t$) + primer ($p$) $\rightarrow$ product ($tp$). The equilibrium constant for the template-primer bimolecular hybridization can thus be expressed as follows:

$$K_{eq} = \frac{c_{tp}}{c_t c_p} \tag{15}$$

We define fractional conversion $\chi$ as the fraction of the reactant that is converted into product, hence expressing in terms of template conversion:

$$\chi_t \equiv \frac{c_{tp}}{c_{0,t}} \tag{16}$$

Consequently, we can express the law of mass action in terms of template fractional conversion and initial single-stranded reactant concentrations, expressing primer concentration according to the bimolecular stoichiometry:

$$K_{eq} = \frac{c_{0,t}\chi_t}{(c_{0,t} - \chi_t c_{0,t})(c_{0,p} - \chi_t c_{0,t})} \tag{17}$$

Since the initial concentrations are normally specified, the equation is only a function of $\chi_t$. Manipulating the equation, we can arrive on the following quadratic equation that gives a solution for $\chi_t$ in terms of $K_{eq}$ and initial concentration values:

$$K_{eq}(c_{0,t} - \chi_t c_{0,t})(c_{0,p} - \chi_t c_{0,t}) = c_{0,t}\chi_t$$

$$K_{eq}(c_{0,t}c_{0,p} - {c_{0,t}}^2\chi_t - c_{0,t}c_{0,p}\chi_t + {c_{0,t}}^2{\chi_t}^2) = c_{0,t}\chi_t$$

$$K_{eq}c_{0,t}{\chi_t}^2 - [1 + K_{eq}(c_{0,t} + c_{0,p})]\,\chi_t + K_{eq}c_{0,p} = 0$$

$${\chi_t}^2 - \left[1 + \frac{c_{0,t} + {K_{eq}}^{-1}}{c_{0,t}}\right]\chi_t + \frac{c_{0,p}}{c_{0,t}} = 0 \tag{19}$$

Expanding $K_{eq} = exp(-\frac{\Delta G^{i,j}_{\text{hyb}}}{RT})$ and solving the resulting equation using the quadratic formula yields:

$$\chi_t = \frac{1}{2}\left[1 + \alpha + \frac{\alpha}{c_{0,p}}\exp\left(\frac{\Delta G^{i,j}_{\text{hyb}}}{RT}\right) - \sqrt{\left(1 + \alpha + \frac{\alpha}{c_{0,p}}\exp\left(\frac{\Delta G^{i,j}_{\text{hyb}}}{RT}\right)\right)^2 - 4\alpha}\,\right] \tag{20}$$

Where:

$$\alpha = \frac{c_{0,p}}{c_{0,t}} \tag{21}$$

The resulting equation expresses the fractional conversion in terms of temperature, $\alpha$, and $\Delta G^{i,j}_{\text{hyb}}$, the latter is in turn a function of salt concentration as well as temperature, hence equation 20 is ultimately dependent only on temperature, salt concentration, and $\alpha$. Despite that in a typical PCR mixture, template concentration in terms of moles is often difficult, if not impossible, to determine, the model still does not collapse for uncertain values of template concentration as is explained in the paper.

We then define fractional specificity $\psi$ as the ratio of target ($T$) to non-target ($N$) fractional conversion:

$$\psi \equiv \frac{\chi_t\big|_{\Delta G^{T,p}_{\text{hyb}}}}{\chi_t\big|_{\Delta G^{N,p}_{\text{hyb}}}} \tag{22}$$

This equation represents the ratio of desirable target product concentration to the undesirable non-target species product concentration. A primer with $\psi$ value of 1000 means that in the presence of a non-target template, the primer will hybridize with such template 1000 times less probable than with a desirable target template. This will result to a loss of amplification signal due to the primer not binding efficiently with the non-target species, a preferable scenario in the design of species-specific primers. This parameter is the primary basis for the screening of candidate target-specific primers among the set of all primer sequences possible for a sequence of a specific genetic marker.

### 4 Modelling Extension Inhibition

In this section, modelling the extension inhibition stated in the third assumption is explained. The phenomenon is similar to the formation of Okazaki fragments in the lagging strand, wherein extension facilitated by the DNA polymerase is halted upon encountering a 5' nucleotide terminal bound to the template. A nick is thus present between the final nucleotide added by the polymerase and the 5' nucleotide terminal. In nature, this is joined together by the ligase enzyme. PCR reaction mixtures do not include the ligase enzyme; hence these nicks cannot be joined together, resulting to two distinct fragments: a fragment extended through the primer that produces a larger amplicon size, and a fragment extended by the primer binding along the former's amplification path. In the next cycle of the amplification though, the first fragment will not be amplified further because it lacks the reverse primer (RP) binding region that is otherwise present in the second fragment. It hence results to the inhibition of product amplification of the first fragment. The whole process is illustrated through a schematic in figure S5. Such an event explains some of the results observed in the laboratory (preliminary in-house experiments) wherein primers amplifying shorter amplicons tend to be highly specific even with poor $\psi$ values while longer amplicons are easily inhibited (fainter gel bands) by non-specific priming.

In modelling this phenomenon, we assume a probabilistic behavior among the binding primers. These probabilities are assumed equivalent to the fractional template conversion $\chi_t$, and are thus deterministic for a given DNA template. The product fraction from a certain primer $i$ is therefore computed by multiplying its initially computed equilibrium fraction $\chi_{0,i}$ to the chain of probabilities from each $j$th primer from the set of primers that may bind along its extension path. Mathematically, it is expressed as:

$$\chi_i = \chi_{0,i} * \prod_{j \neq i} (1 - \chi_j) \tag{23}$$
